# Population dynamics of ribozymes during *in vitro* selection

**DOI:** 10.1101/2025.08.31.673163

**Authors:** Elizabeth M. Brooks, Natalie R. Kotlin, Zoe Weiss, Saurja DasGupta

**Affiliations:** Department of Biological Sciences, University of Notre Dame, Notre Dame, IN 46556, United States; Department of Chemistry and Biochemistry, University of Notre Dame, Notre Dame, IN 46556, United States; Harvard/Massachusetts Institute of Technology MD-PhD Program, Harvard Medical School, Boston, MA 02115, United States; Berthiaume Institute for Precision Health, University of Notre Dame, Notre Dame, IN 46556, United States

## Abstract

Ribozymes are central to models of RNA-based primordial life. Understanding how ribozymes emerge from RNA populations under changing selection pressures can help delineate the biochemical constraints and capabilities of early RNA-based biology. *In vitro* selection has been instrumental in isolating ribozymes with diverse functions from combinatorial populations; however, their population dynamics during selection remain poorly characterized. By analyzing high-throughput sequencing data produced from all rounds of an *in vitro* selection experiment, consisting of over 5 million unique sequences, we tracked how sequences rose and fell in abundance during selection. We found that the most abundant sequences emerged early and maintained their dominance, suggesting that only a few cycles of selection-amplification may be sufficient to isolate well-adapted ribozymes. Characterization of the catalytic activities of the dominant sequences in each selection round indicated that abundance did not always predict activity. This observation was reinforced through nucleotide conservation analysis and activity assays of close sequence variants of the most dominant ribozyme. Our analyses show that sequence complementarity with the substrate emerged as a common feature as early as the first round and persisted throughout, highlighting early convergence of diverse sequence populations on a beneficial catalytic strategy. By combining bioinformatics and experimental approaches, our work presents a detailed description of ribozyme population dynamics during an *in vitro* selection experiment and provides a high-resolution view of how functional RNA sequences compete for survival and propagation under selection pressure.

## INTRODUCTION

Darwinian evolution is a defining feature of life (Benner, 2010). Considering the perceived central role of RNA in carrying genetic information and catalyzing chemical reactions in primordial life, the sustained replication and adaptation of functional RNA molecules would have been key to the origin and evolution of life in an RNA World. Natural selection would have driven diversification of the catalytic capabilities of RNA, leading to the emergence of novel adaptive solutions to the changing environmental conditions on a primordial Earth (Holland & Domingo, 1998; Spiegelman, 1971). *In vitro* selection, which replicates the Darwinian evolutionary process in a test-tube, has enabled the exploration of nucleic acid sequence space and the discovery of novel nucleic acid functions in a laboratory setting (Chen et al., 2007; Jijakli et al., 2016; Robertson & Joyce, 2012; Wilson & Szostak, 1999). Most *in vitro* selections focus on the outputs from the final round; therefore, the details of the evolutionary process that begins with a random collection of sequences and culminates in populations highly enriched in a small subset of those sequences – remain poorly understood. This knowledge gap was recognized in early studies with DNA catalysts and aptamers that investigated the population dynamics of these functional DNA sequences across selection rounds by sequencing individual clones from each round (Schlosser & Li, 2005; Yang et al., 2009). As these early studies preceded the advent of high-throughput sequencing (HTS), only a few hundred sequences could be analyzed in total, yielding an incomplete picture of their evolutionary dynamics. In contrast, the vast sequence coverage allowed by HTS not only enables a more detailed analysis of the selection process but also allows researchers to track its progress across each round. The last decade has seen a rise in the use of HTS to analyze *in vitro* selection experiments, mostly to identify functional sequences isolated in the last round for downstream characterization. Only a handful of studies used HTS to probe the RNA sequence space and even fewer, the evolutionary process. For example, the Ferré D’Amaré (Pitt & Ferre-D’Amare, 2010), Chen (Jimenez et al., 2013; Pressman et al., 2019), Yokobayashi (Kobori et al., 2015), and Hayden (Peri et al., 2022) labs used HTS to generate mutational fitness landscapes of functional RNAs like aptamers and ribozymes. Other studies probed the selection process deeper by focusing on the emergence and conservation of structural motifs in aptamers (Ditzler et al., 2013; Schutze et al., 2011) or broad population-level changes in ribozymes (Ameta et al., 2014; Fu et al., 2012; Pressman et al., 2017) without capturing the fine-grained sequence-level dynamics or the dynamics at single-nucleotide resolution. Aptamer selection has also been studied using computational tools, which have provided useful insight into the design of combinatorial selection experiments (Asif & Orenstein, 2020; Hoinka et al., 2015). Although collectively these findings have improved our understanding of *in vitro* selection and the distribution of functional nucleic acids in sequence space, critical gaps remain in our understanding of how catalytic RNA sequences or ribozymes emerge from large RNA populations and what genetic and structural features contribute to their survival and enrichment. A quantitative picture of the selection dynamics of ribozyme populations during *in vitro* selection will reveal how ribozyme sequences respond to changing selection pressures and consequently, how their relative abundances change over the course of selection.

*In vitro* selection and directed evolution campaigns have paid special attention to ribozyme-catalyzed ligation and polymerization of oligomeric and monomeric RNA building blocks, respectively, as these processes would have been important for replicating RNA genomes and generating RNA enzymes in an RNA World, and consequently, are relevant to the origins of life (Martin et al., 2015). In prior work, we used *in vitro* selection to isolate ribozymes that catalyze the template-dependent ligation of RNA oligomers 5′-activated with the prebiotically relevant 2-aminoimidazole (2AI) group (Walton et al., 2020). As 2AI-activated oligoribonucleotides are also amenable to nonenzymatic assembly, ribozymes that use these building blocks as substrates for enzymatic ligation may represent key intermediates in the transition from nonenzymatic to enzymatic RNA assembly – a major transition in the origin of life. In that work, we used a 5′-2AI-activated, 3′-biotinylated substrate as bait to capture active sequences from an RNA library (∼10^14^ sequences) on streptavidin-coated magnetic beads (Fig. 1A). In addition to bead-based selection of sequences with ligase activity, reverse complementarity of the reverse transcription primer with the newly ligated substrate ensured that only the library sequences that catalyze the desired ligation reaction get selected (Walton et al., 2020). In this work, we employed a combination of standard and custom bioinformatics tools (Supplementary Data Table 1) to analyze HTS data from all eight selection rounds of this ‘AI-ligase’ selection experiment, each containing over a million reads (Supplementary Data Table 2). This approach allowed us to perform a quantitative analysis of the population dynamics of AI-ligase ribozymes at the sequence and nucleotide levels. We tracked the emergence and measured the catalytic activities of the most abundant sequences after each selection round. We followed the rise and fall of RNA sub-populations under increasing selection pressures, which allowed us to observe sequence-level competition for survival. We identified the most conserved nucleotide positions in the close-sequence variants of the most dominant ribozyme and experimentally tested the functional importance of those conserved positions through mutagenesis assays. By tracking ribozyme emergence at different points in the selection, we discovered that at least 88% of all sequences isolated in each round exhibited base-pair complementarity of at least 5 nucleotides with the substrate, highlighting base-pairing as a key catalytic strategy that emerged early and persisted throughout selection. Together, this study provides a multi-layered view of how functional RNAs may respond to selection pressure and reveals the sequence determinants that may drive their fitness and adaptation.

**Figure 1.**
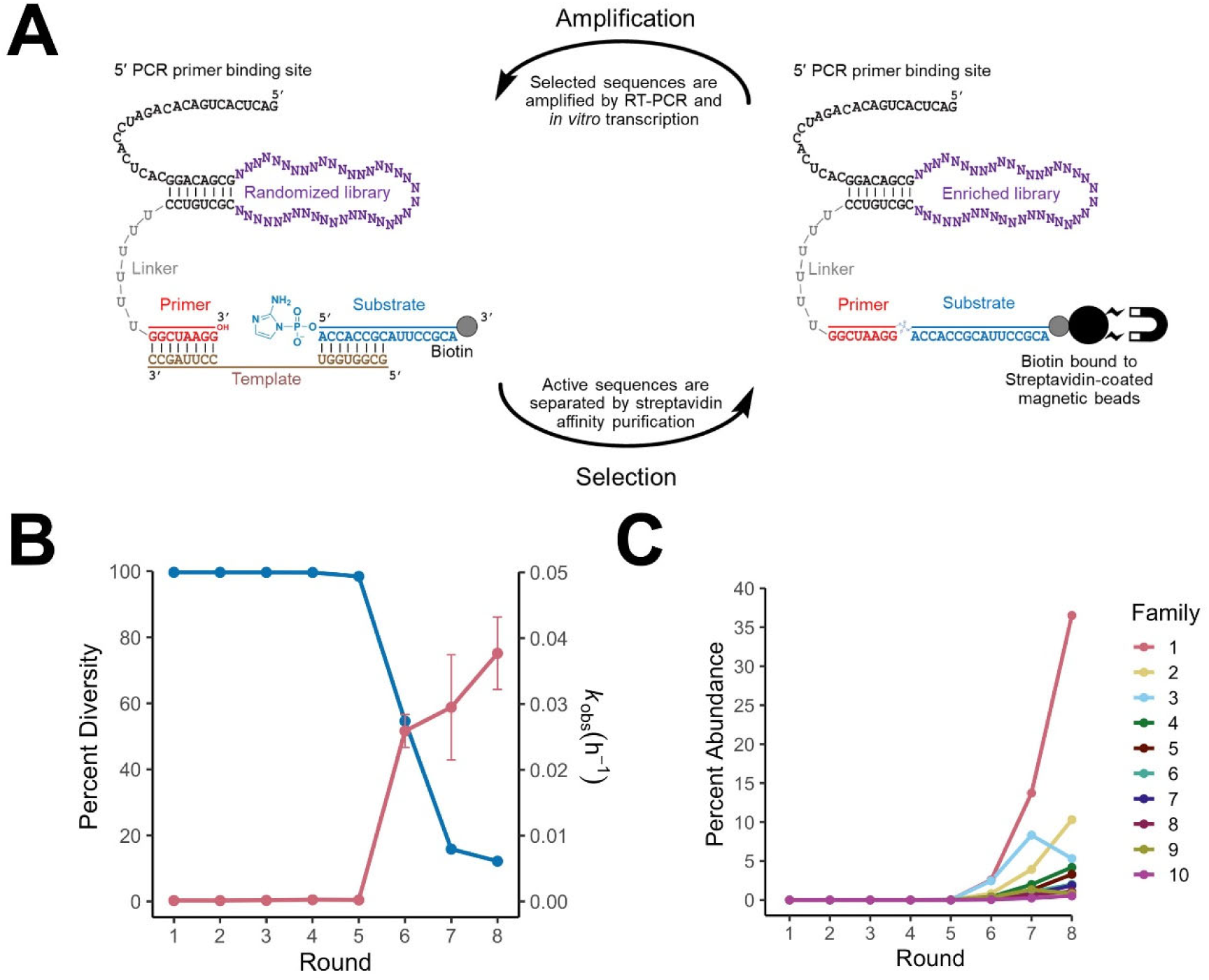
Selection of ribozymes that catalyze the ligation of 2-aminoimidazolide RNA oligonucleotides. **(A)** Ligase ribozyme selection utilized a randomized RNA library with a 40-nucleotide (nt) variable region (purple) flanked by a constant 8 base-pair stem. The 5*’* end of the library contained a 25-nt primer binding site for PCR amplification. The 8-nt primer sequence (red) at the 3*’* end of the library was connected to the catalytic domain by a 6-nt polyuridine linker (gray). A 16-nt template (brown) base pairs with the primer and the first eight nucleotides of the 16-nt 2AI-activated substrate (AI-Lig, blue), leaving an 8-nt overhang on the substrate 3*’* end (referred to as the substrate 3*’* overhang). **(B)** Catalytic activity was detected at round 6, which was accompanied by a marked decrease in sequence diversity (98.44% to 54.61%). This trend was maintained in rounds 7 (15.86%) and 8 (12.20%), which indicated an enrichment in catalytic sequences. Percent sequence diversity is shown in blue and marked on the left y-axis. *k*_obs_ is shown in red and is marked on the right y-axis. **(C)** Sequences contained in the ten most abundant ribozyme families became prominently enriched in round 6, with sequences in Family 1 covering ∼36% of the entire RNA population in round 8.

## RESULTS

### Enrichment of dominant sequences accompanies decrease in sequence diversity

In our earlier study on the selection of AI-ligase ribozymes, we analyzed sequencing data from the final selection round and biochemically characterized only a handful of sequences (Walton et al., 2020). In this work, we employed a bioinformatics pipeline (Supplementary Fig. 1, Supplementary Data Table 1) to analyze sequencing data from all eight rounds of that selection, enabling a quantitative view of the population dynamics of ligase ribozymes under increasing selection pressures. Sequence populations in the early rounds (rounds 1-5), containing an average of ∼992,000 reads per round, remained highly diverse (Supplementary Data Table 2). This modest sequence enrichment in early rounds is consistent with the minimal catalytic activity of the resulting pools (Fig. 1B). Ligase activity became detectable in the RNA library at round 6, which was accompanied by a sharp drop in sequence diversity from ∼98% to ∼55% (Fig. 1B). Enrichment continued through round 8, where the RNA population consisted of only ∼12% unique sequences and catalyzed ligation ∼200-fold faster than the RNA population that emerged at round 1, in which ∼99% of the sequences were unique (Fig. 1B, Supplementary Data Table 2). This marked enrichment in round 6 may have resulted from increased selection pressure from lowering the reaction time from 30 to 10 min (Supplementary Data Table 2). A decrease in the number of unique sequences in round 8 allowed us to cluster similar sequences into sequence families and identify the most abundant sequence in each family, referred to as the ‘peak sequence’ (Supplementary Data Table 3). Given the high degree of sequence similarity within each family and the high level of divergence between families, each family member is assumed to adopt a similar structure and represents a unique class of AI-ligase ribozyme. During the course of selection, sequence families expanded to include single and double mutants of the peak sequences due to error-prone replication by *Taq* polymerase during PCR amplification. This resulted in an increase in sequence diversity within each family, with the most abundant families being the most diverse (Supplementary Fig. 2). Tracking the abundance of each family (a sum of read counts of its member sequences/total read counts) across eight rounds of selection revealed Family 1 as the predominant cluster covering ∼36% of the entire final round population, with its peak sequence (F1 peak) covering ∼79% of that cluster (Supplementary Data Table 3). These dominant sequence families were significantly enriched in round 6 (Fig. 1C), consistent with the decrease in sequence diversity and increase in ligase activity of the RNA population from round 6 (Fig. 1B). The enrichment pattern of sequence families, which considers the relative abundances of all sequences in each family, paralleled that of their peak sequences as revealed in our prior work (Walton et al., 2020).

### Population dynamics at the sequence level

Tracking the relative abundances of the top ten families across eight rounds revealed how they were enriched during selection, but did not capture how the relative abundances of individual sequences in the population changed under the increasing selection pressure. Previous studies that tracked sequence abundance of catalytic DNAs across different rounds of *in vitro* selection showed that the most abundant sequences in earlier rounds can go ‘extinct’ as more active variants emerge (Schlosser & Li, 2005; Yang et al., 2009). This observed extinction could have been due to limited coverage of the sequence space as these studies involved sequencing individual clones for each round, generating only a few hundred sequence reads in total for analysis. By leveraging the larger sequence coverage of HTS, we were able to investigate how the composition of the ‘fittest’ group of ribozymes changed across eight selection rounds (Table 1), each of which contained ∼0.5 to 1 million sequence reads (Supplementary Data Table 2). We counted the number of reads of all unique sequences in rounds 1-8 and identified the ten most abundant sequences in each round, regardless of which round 8 families they belong to (Table 1, Fig. 2). We then tracked these sequences through the other seven rounds to determine how their relative abundances fluctuated along the selection trajectory (Fig. 3), which revealed some general trends in the evolutionary dynamics of RNA, and some specific features of this ribozyme selection.

**Figure 2.**
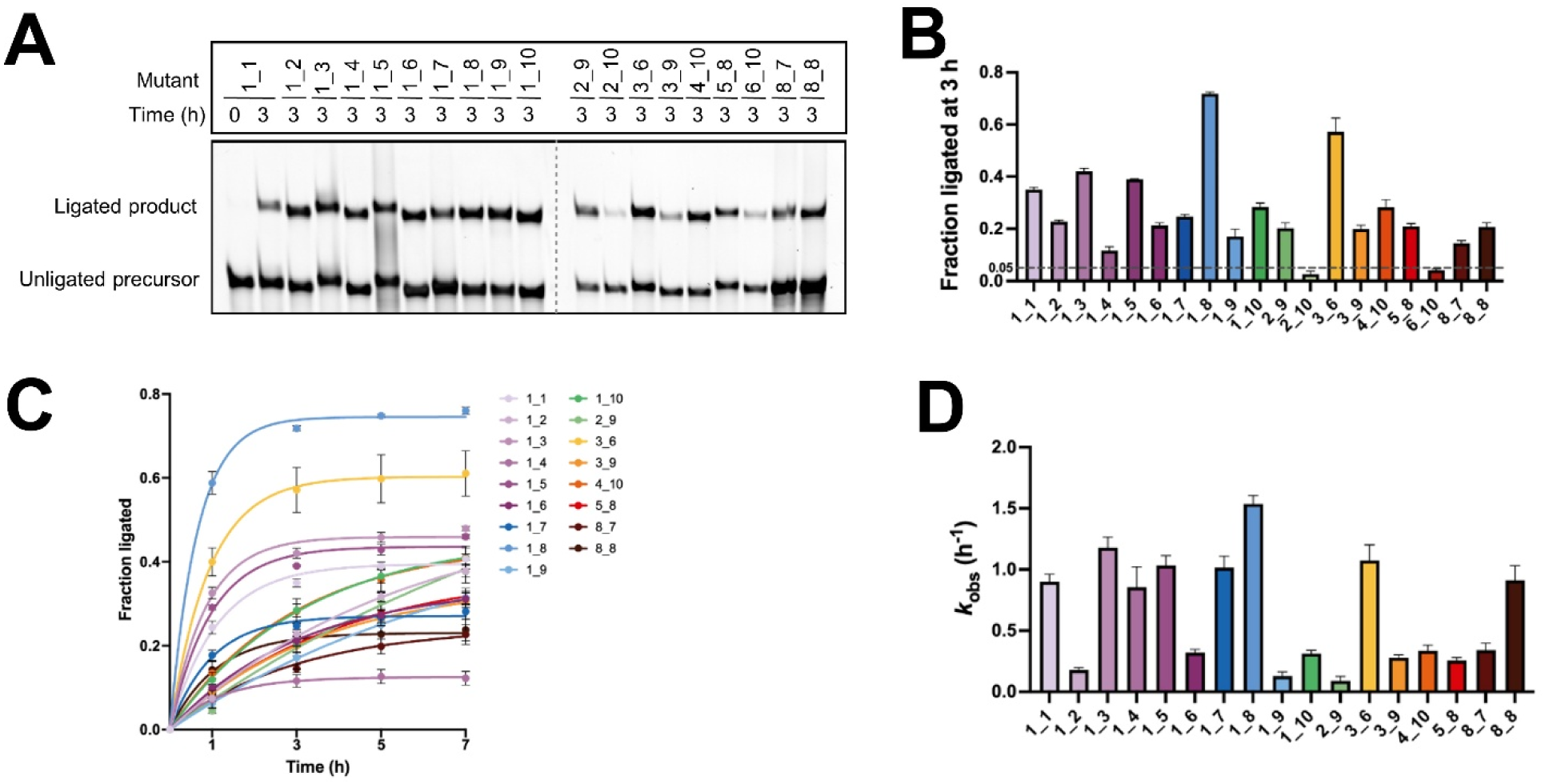
Ligase activity of the nineteen sequences that collectively comprise the ten most abundant sequences in each round (see Table 1, Supplementary data Table 4). **(A)** Ligation assay with the top ten sequences in each round. **(B)** The ten most abundant sequences in each round ligated to different extents in 3 h. **(C)** Kinetic characterization of the sequences in (B) that ligated to at least 5% in 3 h. **(D)** Ligation rate constants for sequences in (C). Error bars in (B) and (C) indicate standard deviation, and in (D) indicate standard error of the mean (S.E.M). All data were obtained from triplicate measurements. Ligation reactions contained 1 μM ribozyme, 1.2 μM RNA template, and 2 μM RNA substrate in 100 mM Tris-HCl (pH 8.0), 300 mM NaCl, and 10 mM MgCl_2_.

**Figure 3.**
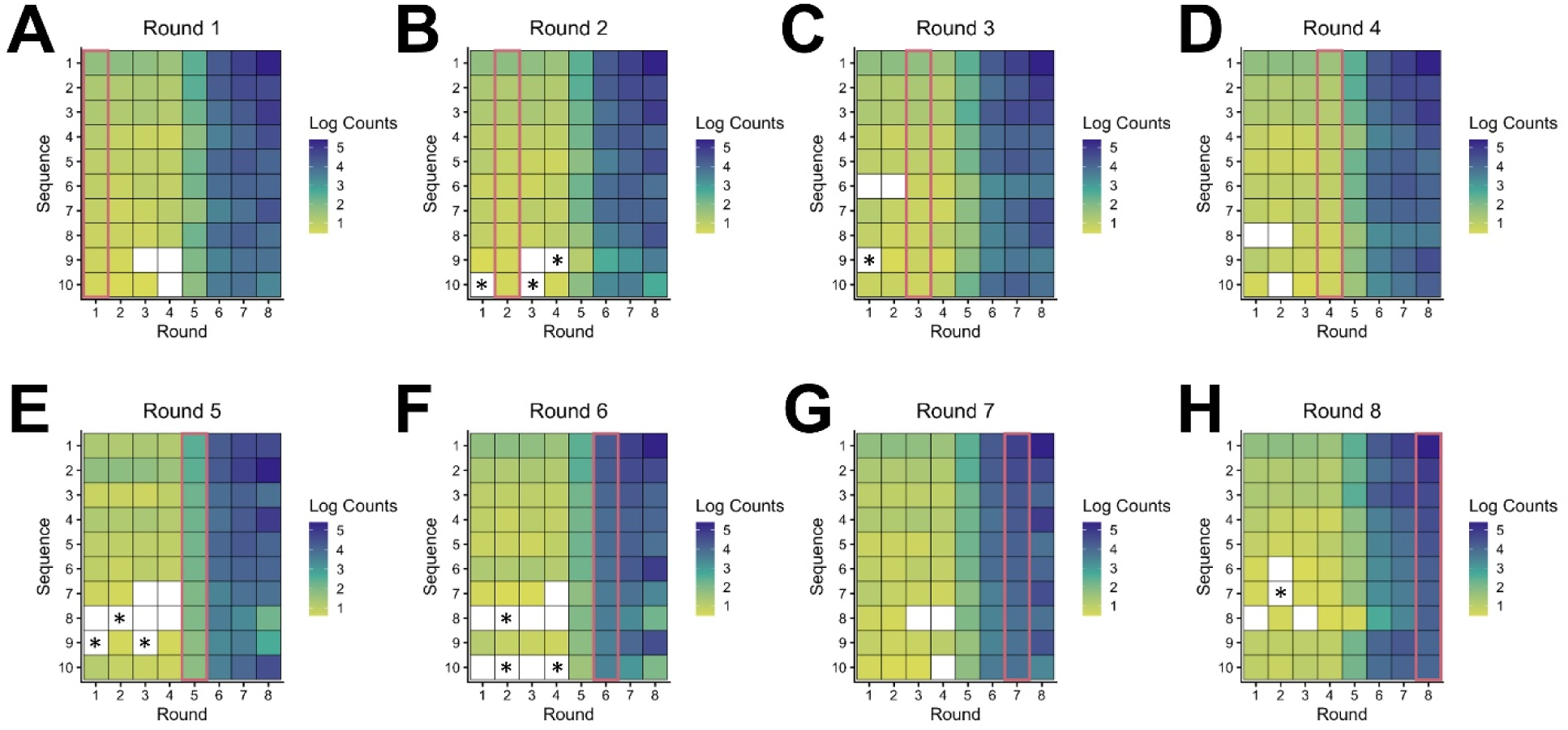
Sequence-level population dynamics of ligase ribozymes during selection. Log_10_ counts of sequencing reads for the ten most abundant sequences in each round plotted across rounds 1 through 8. (**A-H**). For example, in (E), the top 10 sequences in round 5 are tracked back to rounds 1-4 and forward to rounds 6-8, where these sequences are not necessarily the top 10 sequences in the other rounds. Darker blue squares represent higher read counts, lighter yellow squares represent lower read counts, and green squares represent intermediate read counts. The red box outlines the top 10 sequences for that round. White squares indicate read counts below 3. Sequence enrichment was observed beginning in round 5 and prominently in round 6. The most abundant sequences in round 5 survived through to round 8. However, not all of these sequences could be tracked to earlier rounds. Certain sequences fell below 3 reads and even became extinct (read count = 0; marked with a black asterisk) but re-emerged during the course of selection in response to changing selection conditions.

**Table 1.** The composition of the ten most abundant ligase ribozyme sequences changes over the course of selection. Boxes representing the ribozyme sequences are colored according to their ligase activities in 3 h (see Fig. 2A). The eight most abundant sequences from round 1 remain among the top ten through round 4. Of these, the top six sequences from round 1 continue to rank within the top ten throughout all eight rounds, although their relative rankings fluctuate between rounds. BPC indicates the extent of base-pair complementarity with the 8-nt substrate 3’ overhang (see ‘Emergence of substrate complementarity as an adaptive strategy’). Mismatch is indicated by a ‘_’; for example, 3_3 indicates that this sequence (Seq1_8 here) can form 6 base pairs with the substrate 3’ overhang, where two stretches of 3 base-pair interactions are interrupted by a mismatch. Sequences are named according to the round and their rank in the round when they first emerged. For example, Seq1_7 indicates that this sequence emerged first as the 7^th^ most abundant sequence in round 1. See Supplementary Data Table 4 for sequences of the 40-nt variable region of these ribozymes, and Fig. 3 for a detailed illustration of sequence dynamics across selection rounds for all sequences with >2 reads.

|  | Sequence Rank in each round 1-8 |  |  |  |  |  |  |  |  |
| --- | --- | --- | --- | --- | --- | --- | --- | --- | --- |
| Seq ID | 1 | 2 | 3 | 4 | 5 | 6 | 7 | 8 | BPC |
| Seq1_1 | 1 | 1 | 1 | 1 | 2 | 1 | 1 | 1 | 6/8 |
| Seq1_2 | 2 | 2 | 3 | 2 | 1 | 2 | 2 | 3 | 6/8 |
| Seq1_3 | 3 | 3 | 2 | 3 | 4 | 6 | 4 | 2 | 6/8 |
| Seq1_4 | 4 | 5 | 7 | 9 | 10 | 9 | 7 | 4 | 6/8 |
| Seq1_5 | 5 | 4 | 5 | 6 | 5 | 3 | 3 | 9 | 7/8 |
| Seq1_6 | 6 | 7 | 4 | 7 | 6 | 4 | 6 | 10 | 6/8 |
| Seq1_7 | 7 | 8 | 8 | 4 | - | - | 9 | 5 | 8/8 |
| Seq1_8 | 8 | 6 | 10 | 5 | 3 | 5 | 5 | - | (3_3)/8 |
| Seq1_9 | 9 | - | - | - | 7 | - | 8 | - | 5/8 |
| Seq1_10 | 10 | - | - | - | - | 7 | 10 | - | 6/8 |
| Seq2_9 | - | 9 | - | - | - | - | - | - | 5/8 |
| Seq2_10 | - | 10 | - | - | 9 | - | - | - | 5/8 |
| Seq3_6 | - | - | 6 | 8 | - | - | - | - | 6/8 |
| Seq3_9 | - | - | 9 | - | - | - | - | - | (3_4)/8 |
| Seq4_10 | - | - | - | 10 | - | - | - | 6 | 7/8 |
| Seq5_8 | - | - | - | - | 8 | 8 | - | - | (3_4)/8 |
| Seq6_10 | - | - | - | - | - | 10 | - | - | 6/8 |
| Seq8_7 | - | - | - | - | - | - | - | 7 | (3_3)/8 |
| Seq8_8 | - | - | - | - | - | - | - | 8 | 6/8 |

Fraction ligated at 3 h

The most general trend among the top-ranked sequences was the sharp increase in sequence abundance in round 5, followed by another significant increase in round 6 (Fig. 3). This correlates with the repeated changes in selection pressure. In contrast, a constant selection pressure would not promote continued shifts in sequence abundance. Our results show that although population-level activity (Fig. 1B) is not detected in earlier rounds, individual dominant sequences are detected by HTS early on in the selection (Table 1). The ability to detect active sequences before bulk activity in the selection pool underscores the utility of HTS for assessing the potential success of selection experiments at their initial stages. The top six sequences in round 1 maintained their superior fitness and remained in the top ten in subsequent rounds (Table 1, Supplementary Data Table 4). The 7^th^ ranked sequence in round 1 (Seq1_7) fell off the top 10 in rounds 5 and 6, but reemerged in round 8. Seq1_8, on the other hand, maintained high fitness through round 7, but fell out of the top 10 in the final round. In contrast, Seq1_9 and Seq1_10 were only occasionally present in the top 10. Despite these fluctuations, the top 10 sequences in round 1 (before catalytic enrichment) also comprised the top 10 sequences in round 7 (after catalytic enrichment), albeit ranked differently (Table 1, Supplementary Data Table 4), and 7 of the top 10 sequences in round 8 were already abundant in earlier rounds, some as early as round 1. The top 10 sequences in round 8 did not fully align with the peak sequences of the top 10 families (Supplementary Data Table 4). Nine of the ten sequences that did not represent any of the round 8 peak sequences became more abundant in round 7 and fell in round 8, with the exception of Seq8_8, which is one mutation from the F1 peak sequence (Supplementary Fig. 3). Our ability to glimpse the competition among the top sequences in each round to remain dominant allowed us to test the correlation between the abundances of these high-fitness sequences and their catalytic activities. We measured ligation yields for all 19 ribozyme sequences (Seq1_1 to Seq8_8; Table 1, Supplementary Data Table 4, Fig. 2A, B) under identical reaction conditions that included 10 mM Mg^2+^, and characterized ligation kinetics of a subset of sequences that ligated to at least 5% in 3 h. Unsurprisingly, Seq1_1 to Seq1_6, sequences that were dominant in all 8 rounds, ligated to over 5% under these reaction conditions, with Seq1_3 showing the highest and Seq1_4 the lowest yields in this cohort. Interestingly, Seq1_8, which showed the best ligation among all 19 sequences and was the 5^th^ most abundant sequence in round 7, did not make it to the top 10 in the final round, where the Mg^2+^ concentration was reduced from 10 mM to 5 mM, despite its fast kinetics (Table 1, Supplementary Data Table 2, Fig. 2). Another interesting example illustrating the complexity of the selection process is Seq3_9, which emerged only once in the top 10, but was not isolated in the final round. While this sequence showed more efficient ligation (20% in 3 h) compared to another such sequence, Seq6_10 (∼5% in 3 h), its activity was comparable to other sequences such as Seq8_7 and Seq8_8 that emerged for the first time in the final round (Fig. 2).

Merely tracking the composition of the dominant sequences generates an incomplete picture of selection dynamics. Therefore, we tracked the abundances of the ten most abundant sequences in each round across all eight rounds (Fig. 3). Most of the top 10 sequences identified in early rounds persisted throughout selection, although several underwent transient extinction (falling below 3 reads or disappearing entirely) before re-emerging in later rounds. In contrast, many sequences first appeared after round 1 but failed to remain abundant, whereas others emerged only in later rounds and got rapidly enriched. The final-round population contained both long-persisting lineages and newly-emerged sequences, while several previously abundant variants had disappeared. Together, these observations reveal highly dynamic population trajectories during *in vitro* selection, with repeated sequence extinction, re-emergence, and late enrichment, consistent with changing selection pressures and, to a lesser extent, sequence diversification during PCR amplification.

### Population dynamics at the nucleotide level

With an understanding of sequence-level population dynamics of these ligase ribozymes, we sought to understand the linkage between nucleotide enrichment at each position of the isolated ribozyme sequences and their activity. Searching the entire collection of selected sequences for patterns in nucleotide conservation would not yield meaningful information as these sequences belong to different ribozyme families that likely adopt distinct structures. Therefore, we analyzed round 8 sequences with ≥90% sequence similarity (i.e., at least 36/40 nucleotides in the variable region are identical) to the most abundant F1 peak sequence. These closely related variants allowed us to assess how nucleotide identity at each of the 40 variable positions correlated with activity. We quantified the frequency of occurrence of each nucleotide in the 40-nt variable region and observed strong conservation at most positions (Fig. 4A). We assigned an arbitrary conservation threshold of 90%, so that positions 8, 17, 20, and 37, where the predominant nucleotide was <90% abundant, were considered ‘variable’. Conserved positions were expected to be critical to function, while variable positions were expected to be less consequential. We created single-nucleotide mutants at these ‘variable’ positions, as well as at the highly conserved U4 position (∼99% conserved) and tested ligase activity (Fig. 4B-F, Supplementary Fig. 4). Although these mutations occurred in conjunction with other mutations in a subset of the isolated sequences, the majority of sequences with mutations at these positions had no other accompanying mutations, supporting the validity of our mutagenesis experiments.

**Figure 4.**
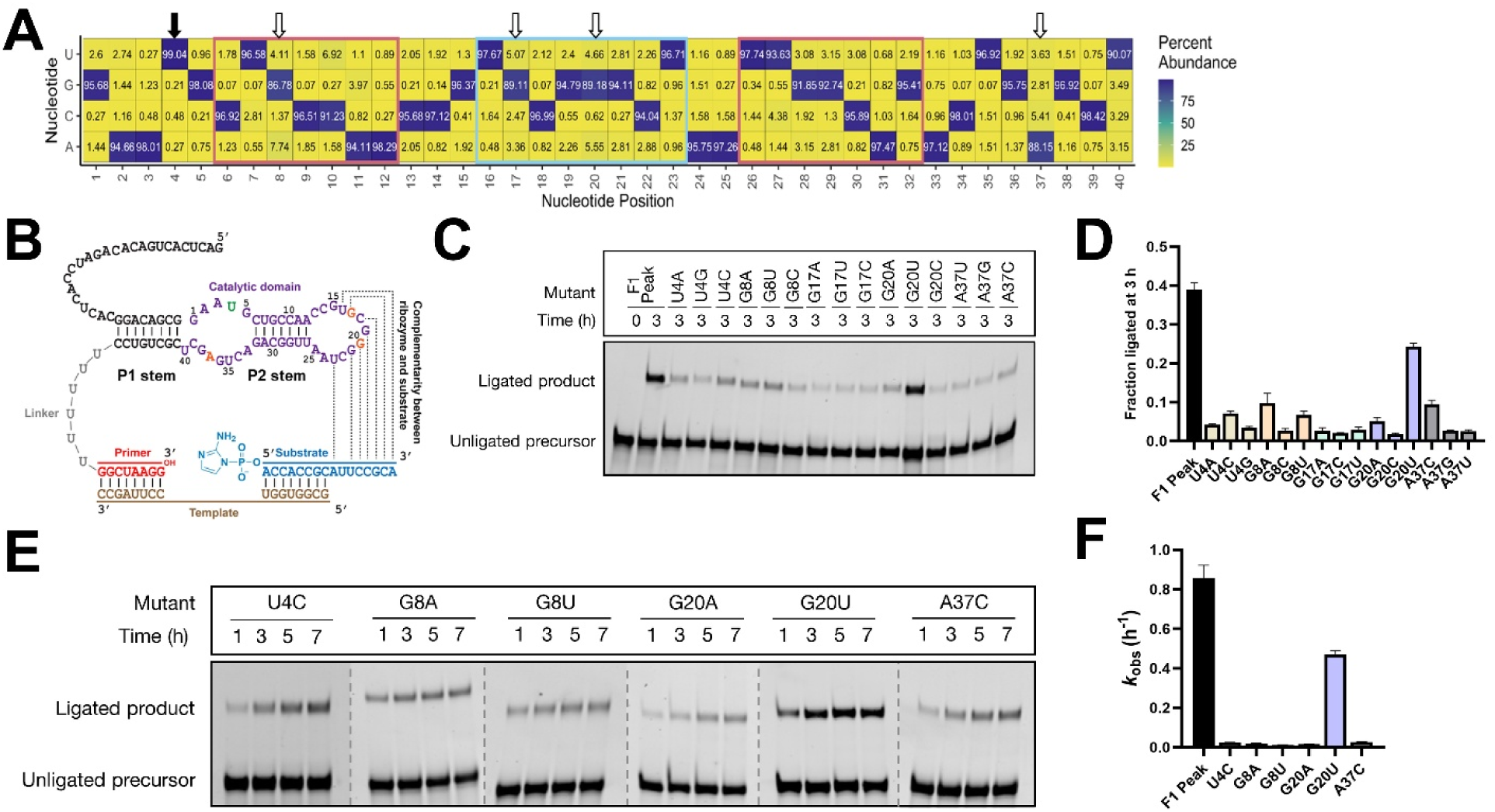
Nucleotide conservation in sequences with ≥90% overlap with the most abundant sequence (F1 peak). **(A)** Fractional abundance of each nucleotide in positions 1-40 of the 40-nt variable region. All 40 positions were moderately to highly conserved. The most variable (position 8) and conserved (position 4, indicated by a black arrow) positions showed ∼86% and ∼99% conservation, respectively, for the predominant nucleotide. Considering our arbitrary threshold of 90% nucleotide conservation, we assigned positions 8, 17, 20, and 37 as ‘variable’ (indicated by white arrows). Nucleotides that comprise base-paired stem P2 of the Family 1 peak sequence are shown in red boxes, and the region complementary to the substrate 3*’* overhang is highlighted in a light-blue box (see Fig. 5A). The percent abundances of each nucleotide were calculated by dividing the number of unique sequences with a given nucleotide by the total number of unique sequences. **(B)** SHAPE-derived secondary structure of the most abundant ligase sequence (F1 peak sequence) bound to the template and substrate. Dashed lines indicate base-pair complementarity between the ribozyme and substrate sequences (Walton et al., 2020). The 40-nt variable region of the ribozyme is depicted in purple. The most conserved nucleotide position (nucleotide 4) is shown in green, and the least conserved positions (8, 17, 20, 37) are shown in red. **(C)** Ligation assay with sequence variants of the F1 peak sequence containing point mutations at the positions indicated in green (4) and red (nucleotides 8, 17, 20, and 37). (**D**) Ligation yields after 3 h for the point mutants of the F1 peak sequence. **(E)** Kinetics gels for single mutants of the F1 peak sequence that ligate to >5% in 3 h: U4C, G8A, G8U, G20A, G20U, and A37C. **(F)** Ligation rate constants for the F1 peak point mutants in (E). Data in (D) and (F) were obtained from triplicate measurements. Error bars in (D) indicate standard deviation, and in (F) indicate standard error of the mean (S.E.M). Ligation reactions contained 1 μM ribozyme, 1.2 μM RNA template, and 2 μM RNA substrate in 100 mM Tris-HCl, pH 8.0, 300 mM NaCl, and 10 mM MgCl_2_.

Of the U4 mutants, only U4C showed detectable ligation at ∼7% in 3 h, supporting the high level of conservation at this position. G8, where the wild-type residue was present in ∼86% of the sequences, was more amenable to mutation, but none of the mutants ligated to more than 10%. G17 was the most resistant to mutation, with none of its variants retaining function. G20U, on the other hand, showed a significant ligation yield of ∼26%, with only a 2-fold lower *k*_obs_ compared to the F1 peak (Fig. 4C-F). A37C was the only A37 mutant to show detectable activity (Fig. 4C, D). Taken together, our mutational analyses show that the degree of nucleotide conservation does not necessarily translate to trends in activity. For example, while the most abundant non-wild-type nucleotide at positions G8 and A37 (A and C, respectively) showed the best ligation yields, this pattern did not hold for G20, where the second most abundant mutant, G20U, ligated most efficiently. We also found that abundance was not a reliable proxy for fitness, as the inactive mutants survived selection, indicating their fitness (Komarova & Kuznetsov, 2019; Martin et al., 2015; Pressman et al., 2019). However, more broadly, the general inefficiency of ligation shown by these F1 peak mutants is consistent with their low abundance in the selected population. Considering that there are apparently very few high-fitness F1 mutants, the fitness peak for this sequence family would most likely be narrow and isolated. Broadening the coverage of the initial sequence library could perhaps lead to the identification of multiple peak sequences as observed by Chen and co-workers (Jimenez et al., 2013; Pressman et al., 2019). The structure of fitness landscapes is expected to be significantly impacted by changes in the environment, where favorable conditions such as molecular crowding may render the landscape less rugged (DasGupta, 2020; DasGupta et al., 2023; Saha et al., 2014).

### Emergence of substrate complementarity as an adaptive strategy

Beyond strong nucleotide conservation in the most abundant ribozyme sequence, we observed that the peak sequences of the ten most abundant families contained a stretch of 5–8 nucleotides partially or fully complementary to the last 8 nucleotides of the substrate (Table 2). This region, referred to as the ‘substrate 3′ overhang’ does not pair with the template during ligation (Fig. 4B, Supplementary Fig. 5A), and is therefore available for base-pairing with the ribozyme. Interactions between the ribozyme and the substrate 3′ overhang presumably help in substrate binding and positioning for the ligation reaction, making it critical for catalysis. This is supported by our earlier work, where deleting this sequence from the substrate or disrupting base-pairing between the substrate and the ribozyme (F1 peak) caused a significant ligation defect (Walton et al., 2020). While most peak sequences possess a stretch of at least 5-nucleotide (nt) complementary to the substrate 3′ overhang, the F7 peak has a single mismatch interrupting this complementary stretch of nucleotides, which still preserves 3-nt complementarity on both sides, making interactions with the substrate viable. F1, F3, F7, and F10 peak sequences exhibit potential for wobble pairing in addition to standard Watson-Crick pairing.

**Table 2.**
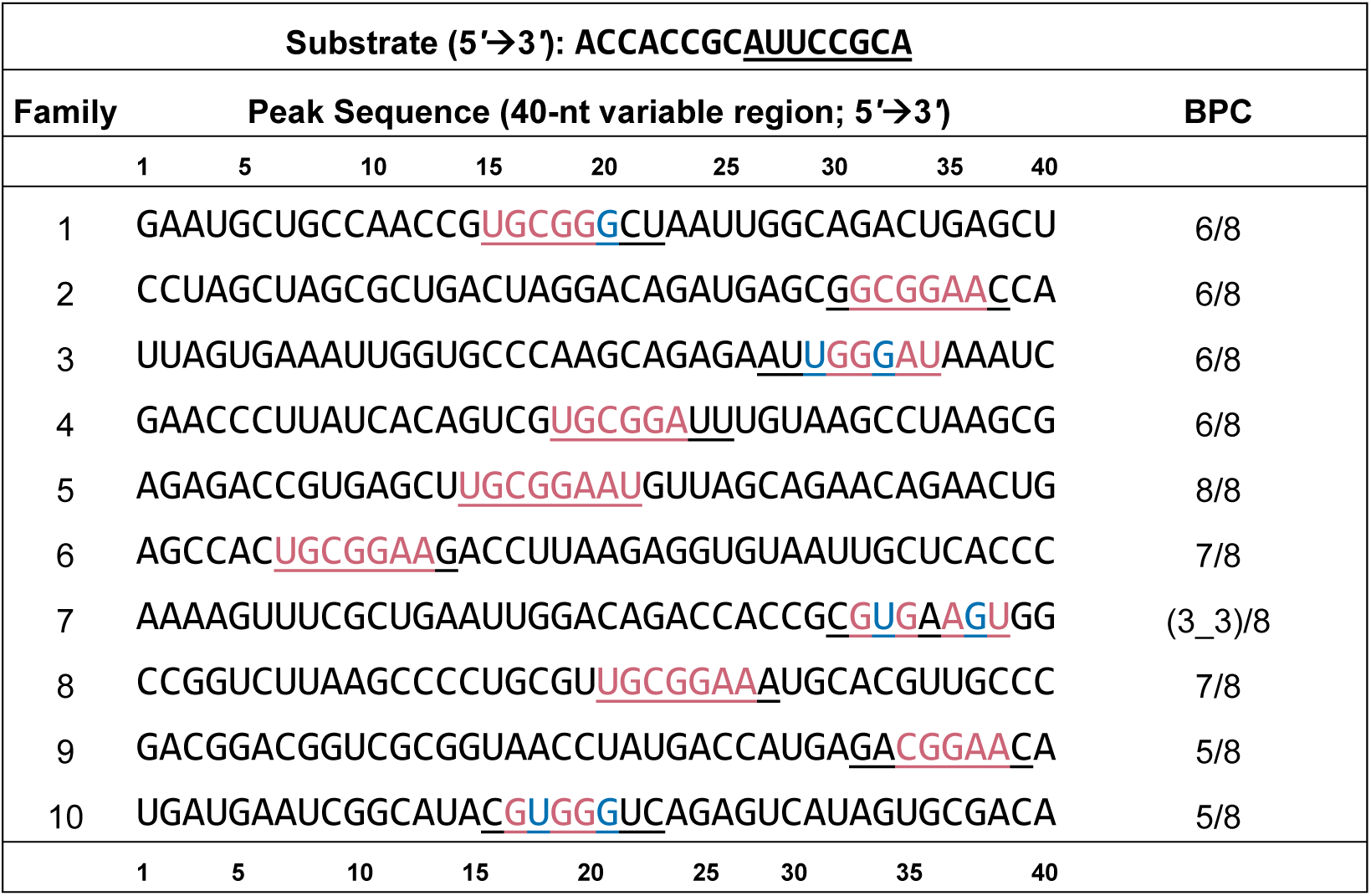
The peak sequences of the ten most abundant families in round 8 have significant base-pair complementarity (BPC) with the substrate sequence. Watson-Crick complementarity with the substrate is indicated in red and wobble complementarity is indicated in blue. The nucleotides contained in the 8-nt sliding window are underlined in each sequence (see Supplementary Fig. 5 and Materials and Methods for details). The extent of complementarity between the peak sequences and the 8-nt substrate 3*’* overhang (underlined) ranges from 5/8 (62.5%) to 8/8 (100%). Peak sequence 7 has two regions of 3-nt complementarity interrupted by a single mismatch, which is indicated by a (3_3)/8 complementarity (see Fig. 5 for a detailed illustration of how sequences with substrate complementarity emerged during selection).

As base pair complementarity between the substrate and ribozyme was found to be a general feature in the most abundant ribozyme families isolated in round 8 (Table 2, Supplementary Fig. 6) and throughout selection (Supplementary Data Table 4), we asked if substrate complementarity contributed to the fitness of the isolated ribozymes. To test this idea, we analyzed sequence data from all eight rounds (∼5.5 million sequence reads) to track the emergence of sequences with at least 5-nt complementarity with the substrate 3′ overhang. Our search algorithm does not distinguish between Watson-Crick and wobble base-pairing and allows up to two internal base-pair mismatches, as long as a complementarity of at least 3-nt is maintained on either side of the mismatch, reflecting the potential for the formation of a moderately stable stem (Supplementary Fig. 5) (see ‘Materials and Methods’).

In a random space of 40-nt sequences, only ∼45% of sequences have ≥5-nt base-pair complementarity with the substrate 3′ overhang, with this fraction dropping to 9.9%, 2.5%, and 0.4% for 6-nt, 7-nt, and 8-nt complementarity, respectively (Fig. 5A). All sequences in this round 0 population have at least 3-nt complementarity to the substrate overhang (considering wobble pairing). Interestingly, the fraction of sequences with ≥5-nt complementarity rose to ∼94% after a single round of selection, with ∼44% exhibiting 6-nt complementarity, a feature that never dropped below 24%. The fraction of sequences with 7-nt complementarity rose to ∼17% in round 1 and never fell below 10%, and sequences with 8-nt complementarity rose to ∼6% in round 1 and to ∼9% in round 8. The amplification of sequences with significant substrate complementarity in round 1 conforms with the emergence of the most fit sequences early in selection (Table 1). The overall abundance of sequences with significant substrate complementarity markedly increased in round 6 (Fig. 5B), consistent with the observed sequence and catalytic enrichment in that round (Figs. 1B, C). Overall, sequence reads with 6-nt complementarity continued to increase through rounds 7 and 8 (Fig. 5B), which is also in agreement with the gradual increase in the fraction of unique sequences with 6-nt complementarity over rounds 6-8 (∼24% in round 6 to ∼40% in round 8) (Fig. 5A).

**Figure 5.**
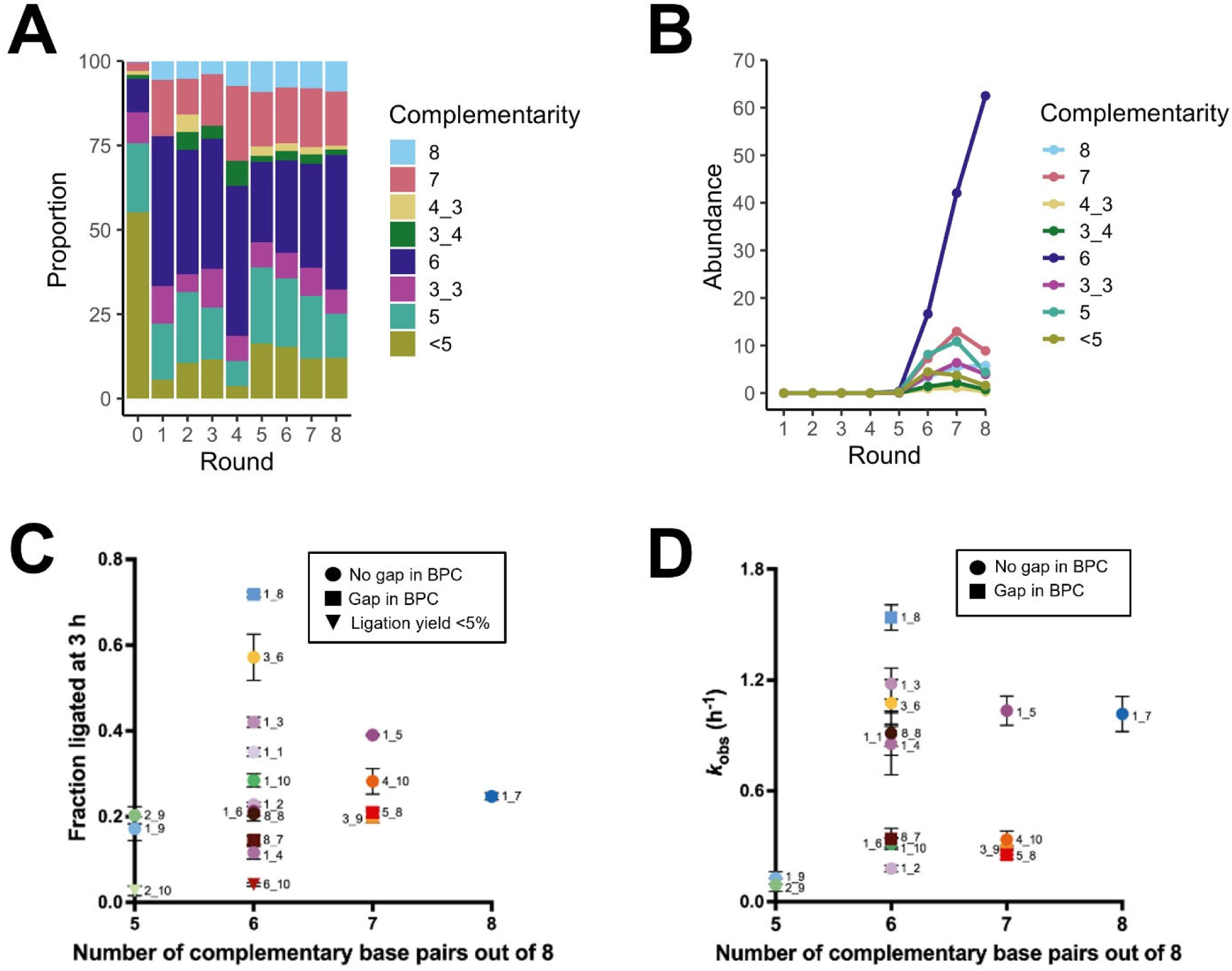
Emergence of ribozymes exhibiting sequence complementarity with the substrate 3′ overhang.#. **(A)** The proportion of complementary nucleotides in the unique sequences (with >2 reads) isolated from all eight rounds of selection and in the round 0 population. The majority (∼90%) of the selected sequences in rounds 1-8 exhibit complementarity of at least 5 nucleotides with the substrate. The abundance of sequences with 6-nt complementarity increased after round 5 from ∼24% to ∼40% (in round 8). Sequences with 8-nt complementarity were enriched beginning in round 4. Sequences with internal mismatches that create gaps are denoted with an underscore (e.g., 3_3 indicates two stretches of 3-nt complementarity separated by a mismatch). **(B)** Sequences with base-pair complementarity with the substrate, including sequences that have mismatches with the substrate 3′ overhang (3_3, 3_4, or 4_3), were enriched in round 6. Sequences with 6-nt complementarity dominated the isolated RNA population through round 8. **(C)** Distribution of sequences that collectively comprise the ten most abundant sequences isolated from each round (nineteen in total) according to their ligation yields in 3 h (see Table 1, Supplementary Data Table 4). Sequences with 5 base-pair complementarity with the substrate 3′ overhang ligated modestly (at or under 30%), but those with >6 base-pair complementary generally performed better. Sequences with 6 base-pair complementarity exhibited a large spread of ligation yields ranging from ∼5% to ∼70%. **(D)** Distribution of sequences from (C) that ligated to at least 5% in 3 h according to their ligation rate constants. All sequences with a base-pair complementarity of 5 ligated slowly, and the single sequence with a base-pair complementarity of 8 exhibited a moderately high *k*_obs_ value. Sequences with 6 base-pair complementarity exhibited a large spread of *k*_obs_ values.

Although the top 10 sequences in each of the eight rounds exhibited base-pair complementarity of at least 5 (Supplementary Data Table 4) bolstering the link between complementarity and ligase fitness, we found that the extent of this complementarity was not directly correlated to catalytic activity. Going back to the ten most abundant sequences from each of the eight selection rounds, we found that consistent with population-level data, most sequences (11/19) had 6-nt base-pair complementarity (BPC) with the substrate (Supplementary Data Table 4, Figs. 5C, D). However, these include sequences with the lowest and highest activities (∼5% ligation by Seq6_10 to ∼70% ligation by Seq1_8 in 3 h). This spread of ligase activity demonstrates a lack of clear correlation between the extent of complementarity and catalytic activity. A caveat here is that all ligation assays were performed at 10 mM Mg^2+^, which is the condition used in selection round 7 and in the characterization of the peak sequences in our earlier work (Walton et al., 2020). Therefore, at higher [Mg^2+^] used in earlier rounds, substrate complementarity could have contributed to ribozyme fitness and driven their selection. Despite the weak correlation between the extent of substrate complementarity and ligase activity, our results suggest that complementarity improves overall ribozyme fitness, even under the weaker selection pressures of the earlier rounds. Sometimes, *in vitro* selection is redirected toward a different target by starting with RNA pools before they are significantly enriched in active sequences. However, in this case, because >90% of the sequences that emerged after the first round already exhibited significant substrate complementarity, it may be difficult to evolve away this substrate sequence dependency from early RNA pools. The overwhelming success of the sequences that possessed substrate complementarity indicates that this ribozyme selection was guided by convergence on this common catalytic strategy.

## DISCUSSION

Historically, the primary purpose of *in vitro* selection has been to isolate functional biomolecules. Therefore, these experiments are generally characterized at their endpoints, and consequently, the dynamics of a large collection of functional sequences competing for survival that ultimately result in the isolation of active molecules remain obscure. To understand how phenotypic selection occurs at the level of RNA genotypes, we analyzed HTS data from all eight rounds of a ligase ribozyme selection experiment. The large coverage provided by HTS allowed us to track the emergence of the dominant sequence families to previous rounds, trace the rise and fall of the most abundant sequences in each round, map nucleotide conservation for the most abundant sequence family, and identify base pair complementarity between isolated ribozymes and the substrate as a convergent catalytic strategy.

Our analysis reveals a complex picture of RNA population dynamics during *in vitro* selection. While the most dominant ribozyme sequences emerged early in the selection process, some initially abundant sequences became undetectable in intermediate rounds before reappearing later, whereas others emerged only during the second half of the selection. These observations indicate that sequences that ultimately dominate the final population are not necessarily the fittest throughout the entire selection trajectory, likely reflecting shifts in selection pressure. This view is further supported by kinetic characterization of the ten most abundant sequences from each of the eight rounds of selection, which showed that some sequences enriched in earlier rounds, but not among the most abundant at the end of selection, exhibited faster ligation kinetics than several of the final dominant sequences. Together, these findings suggest that sequence abundance and catalytic performance are related but not always tightly coupled during evolutionary enrichment (Martin et al., 2015), and underscore the importance of analyzing intermediate rounds to capture transient, yet potentially functional ribozymes. This is also supported by our nucleotide conservation analysis, which showed that the degree of conservation at each position does not always correlate with mutational robustness, and that abundance may not be a suitable proxy for activity. It is possible that changes in the sequence abundances were affected by different processes, such as amplification bias or genetic drift. Genetic drift can result in the stochastic loss or gain of high-fitness rare sequence variants between rounds of selection. Our analysis uncovered a preponderance of ribozymes possessing regions complementary to the substrate, with the extent of complementarity increasing through the course of selection. This highlights the contribution of base-pairing between ribozymes and the substrate toward overall fitness, which is consistent with the inefficiency of ligation using substrates that eliminate this complementarity (Walton et al., 2020). The isolated ligases appear to have converged on substrate complementarity as a key catalytic strategy. However, the extent of complementarity did not correlate strongly to ligase activity.

Although the use of HTS in *in vitro* selection/evolution experiments is not new, previous studies focused on the emergence and conservation of structural motifs or broad population-level changes in aptamers or ribozymes, without capturing the fine-grained sequence-level dynamics or the selection dynamics at single-nucleotide resolution described in this work. The only studies that investigated nucleic acid population dynamics during *in vitro* selection were performed more than two decades ago before the invention of HTS. As a result, this study relied on low-throughput cloning-based sequencing of only a few hundred RNA-cleaving DNAzyme sequences across 17 rounds (Schlosser & Li, 2005). Since then, mathematical models have attempted to simulate population dynamics during *in vitro* selection and evolution, but the simplifying assumptions in these models have limited their ability to reflect the stochastic nature inherent in experimental systems (Higgs & Muller, 2025; Hoinka et al., 2015). In contrast, our study provides an experimentally grounded, high-resolution analysis by simultaneously tracking the abundance and activity of individual sequences during ligase ribozyme selection. This approach enables a more detailed understanding of population dynamics and provides a generalizable framework for dissecting the evolutionary dynamics of functional RNAs.

## MATERIALS AND METHODS

### Materials

All chemical reagents were purchased from Sigma, unless otherwise specified. The hydrochloride salt of 1-ethyl-3-(3-dimethylaminopropyl) carbodiimide was purchased from Alfa Aesar. The hydrochloride salt of 2-aminoimidazole was purchased from Combi-Blocks, Inc. Enzymes were purchased from New England Biolabs unless mentioned. SYBR Gold Nucleic Acid Gel Stain was purchased from ThermoFisher Scientific. 100% ethanol was purchased from Decon Laboratories, Inc. QIAquick PCR purification kits were purchased from Qiagen. The Sequagel-UreaGel concentrate and diluent system for denaturing polyacrylamide gels was purchased from National Diagnostics. Oligonucleotides used in this work are listed in Supplementary Data Table 5 and were purchased from Integrated DNA Technologies (IDT), or generated by *in vitro* transcription of corresponding DNA templates, except NEBNext SR and NEBNext Index proprietary primers for Illumina sequencing were purchased from New England Biolabs. MiSeq Reagent Kit v3, 150 cycles was purchased from Illumina.

### Ribozyme selection and sequencing

The *in vitro* selection protocol is described in detail in the prior study that identified 2-aminoimidazole ligase ribozymes(Walton et al., 2020). In brief, 1 nmol of the RNA library containing a 40-nt random region was incubated with 1.1 and 1.2 equivalents of an RNA template and a 5′ 2AI-activated, 3′-biotinylated substrate, respectively, for 10-120 min, in the presence of 250 mM NaCl, 50 mM Tris-HCl (pH 8), and 5-20 mM MgCl_2_ at room temperature (Supplementary Data Table 1). The reaction was quenched with 200 mM EDTA, 200 µM DNA oligo RQ1 (complementary to the substrate sequence), and 20 µM DNA oligo RQ2 (complementary to the 3′ end of the library). Ligated RNAs were separated from unligated ones by streptavidin bead capture (1 mg MyOne Streptavidin C1 Dynabeads per 100 pmol library), reverse transcribed using Protoscript II, and PCR amplified with *Taq* polymerase to generate dsDNA templates for *in vitro* transcription. RNA obtained from transcription was used as the input library for the next round of selection. NEBNext SR Primer and NEBNext Index Primers for Illumina sequencing were appended to the dsDNA libraries using sequential PCRs using primers, SeqPCR1 and SeqPCR2 for the first PCR and SeqPCR3 and SeqPCR4 for the second PCR. The resultant construct was purified by a Qiagen PCR cleanup kit followed by preparative agarose gel electrophoresis, and sequenced by MiSeq (Illumina, MiSeq Reagent Kit v3, 150 cycles).

### Bioinformatics analysis of high-throughput sequencing data

**1. Overview of the bioinformatics pipeline.** The demultiplexed raw sequencing reads from each round were processed with a standard approach and analyzed using publicly available software. The conventional bioinformatics pipeline was employed using custom BASH and R scripts corresponding to each step of the following analysis. The analysis scripts are available on GitHub (https://github.com/dasguptalab/ligase_ribozyme_population_dynamics) and Zenodo (https://zenodo.org/records/21976743). Sequencing data is available at CurateND (https://doi.org/10.7274/31048348). The BASH (v3.2.57(1)-release) scripts were used to process the files of sequence data from each sequencing round. R (v4.3.2) scripts were created to parse nucleotides within sequences, generate figures, and tabulate results (Team, 2023). The stringr (v1.5.1) (H. Wickham, 2023) R package was used to parse individual nucleotides of sequences and dplyr (v1.1.4) (H. F. Wickham, R.; Henry, L.; Müller, K.; Vaughan, D., 2023) was used to format data tables. The rcartocolor (v 2.1.1) (Nowosad, 2018), scales (v 1.3.0) (Wickham, 2025), and ComplexHeatmap (v 2.16.0) (Gu, 2022) packages were used for visualizing results.
**2. Data pre-processing.** The following steps were performed to filter and prepare the raw data for analysis (Supplementary Fig. 1, Supplementary Data Table 1). First, the raw paired-end reads were merged using FLASH (v1.2.11) (Magoc & Salzberg, 2011) to conserve the most reads (Step 1) (Papastavrou et al., 2024). FLASH also produces files containing paired-end reads for which no good overlap was found, which can be used as a normal paired-end fragment library (Magoc & Salzberg, 2011). Sequencing reads were filtered by an average Phred score of 30 (Q>30) and adapter content was removed using Trimmomatic (v0.39) (Bolger et al., 2014) (Step 2). Sequences that did not contain the predefined 8-nt regions immediately upstream and downstream of the 40-nt variable region were filtered out (Step 3). The filtered sequences were trimmed to the 40-nt variable region by removing these predefined 8-nt regions (Step 4). Next, the merged sequence files were combined with the unmerged read files to retain as much high-quality data as possible (Step 5). Since the sequencing read data contains copies of some sequences, the sequences were de-replicated to keep only the unique sequences and their read counts in each round (Step 6). The de-replicated sequences were filtered to only keep sequences that had >2 reads as they were less likely to be sequencing artefacts (Step 7) (Ameta et al., 2014). The resulting sequences were ordered by abundance in each round according to their read counts (Step 8). Between 92,366 and 1,063,996 unique sequences remained per round after pre-processing. Percent sequence diversity was calculated by dividing the number of unique reads by the number of high-quality reads per round.
**3. Identification and characterization of sequence families.** The unique sequences from the last round of selection (round 8) were clustered using CD-HIT-EST (v4.8.1) (Fu et al., 2012), where each cluster comprised sequences that had ≥90% similarity (sequence identity threshold of 0.9) to each other and consequently, differed by at most 4-nucleotide (nt) positions (i.e., at least 36/40-nt similarity) (Step 9). The 10 most abundant sequence clusters (‘families’) were determined (Step 10). Clustering enabled the detection of representative peak sequences for each ribozyme family (Table 2, Supplementary Data Table 3). The peak sequences from each family were verified to have had at least 10% dissimilarity, confirming that we had identified distinct sequence families. Next, we counted the total reads for each sequence family across all rounds by identifying the sequences with at least 90% similarity to the peak sequence of each family (Supplementary Data Table 2) (Step 11). The percent abundance in each round was calculated by dividing the number of reads for each family by the total number of high-quality reads in that round.
**4. Characterization of population dynamics.** We quantified the abundance of each unique sequence across all 8 rounds of selection (Step 12). Then, we identified the ten most abundant sequences in each round, regardless of their affiliation to sequence families (Supplementary Data Table 4), and represented their abundances in terms of the log-transformed counts of their sequencing reads (Fig. 3) (Step 13). This allowed us to investigate the evolutionary dynamics of the most abundant ribozyme sequences in each round. The percent abundances were calculated by dividing the number of reads for each sequence by the total number of high-quality reads for each round.
**5. Identification of conserved features**. To investigate the extent of nucleotide conservation in the most abundant class of ligase ribozymes (Family 1), we developed a script that calculates the frequency of occurrence of each nucleotide at each of the 40 variable positions (Fig. 4A) (Step 14). The percent abundances of each nucleotide were calculated by dividing the number of unique sequences with a given nucleotide by the total number of unique sequences in the family. This revealed the importance of nucleotide identity at each position for ligase activity of the isolated ribozymes.

To investigate the role of base-pairing between the ribozyme and the substrate, we developed a custom algorithm to determine the extent of complementarity of all sequences isolated during selection (rounds 1-8) with the last eight nucleotides of the substrate (AUUCCGCA) by parsing the individual nucleotides in an 8-nt sliding window. Our algorithm searches for regions in the selected sequences that best match the expected complement sequence (TGCGGAAT) (Supplementary Fig. 5). Wobble pairs were also included in this search. Our algorithm allows for up to two mismatches that create gaps in the complement sequence, as long as there is at least a 3-nt consecutive match, sufficient to form a moderately stable stem. For each round of selection, we calculated the fraction of unique sequences with at least 3-nt complementarity to the substrate overhang (Fig. 5A), irrespective of their read counts (Step 15). A simulated hypothetical round 0 pool of random 40-nt sequences (Fig. 5A) was created using the Linux RANDOM function in a custom BASH script. The correlation between complementarity and fitness was explored by tracking changes in the relative abundances of these sequences (Figs. 5A, B). We also identified conserved complementary regions in the representative peak sequences for the ten most abundant sequence families (Table 2) and all sequences in the ten most abundant families (Supplementary Fig. 6) (Step 16). Additionally, we identified conserved complementary regions in the ten most abundant sequences in each round of selection (Supplementary Data Table 4) (Step 17).

### RNA preparation and ligation assays

Ligase ribozymes were prepared by *in vitro* transcription of corresponding dsDNA templates containing 5′ terminal 2′-*O*-methyl modifications to reduce transcriptional heterogeneity at the 3′ end of the synthesized RNA generated by PCR (Kao et al., 2001). *In vitro* transcription reactions contained 40 mM Tris-HCl (pH 8), 2 mM spermidine, 10 mM NaCl, 25 mM MgCl_2_, 10 mM Dithiothreitol (DTT), 30 U/mL RNase inhibitor murine (NEB), 2.5 U/mL thermostable inorganic pyrophosphatase (TIPPase) (NEB), 4 mM of each NTP, 30 pmol/mL DNA template, and T7 RNA Polymerase (NEB) for 3 h at 37°C. Transcription was quenched with DNase I, and RNA was extracted with phenol-chloroform-isoamyl alcohol (PCI) and purified by denaturing PAGE.

The 2-aminoimidazole-activated substrate was obtained by incubating the corresponding 5′ monophosphorylated oligonucleotide with 0.2 M 1-ethyl-3-(3-dimethylaminopropyl) carbodiimide (HCl salt) and 0.6 M 2-aminoimidazole (HCl salt, pH adjusted to 6) for 2 h at room temperature. The reaction was desalted by washing with water in Amicon Ultra spin columns (3 kDa cutoff) and purified by reverse-phase analytical HPLC using a gradient of 98% to 75% 20 mM TEAB (triethylammonium bicarbonate, pH 8) versus acetonitrile over 40 min.

Ligation reactions (5 μL) contained 5 pmol RNA, 6 pmol RNA template, 10 pmol 2-AI-activated RNA substrate, 300 mM NaCl, 10 mM MgCl_2_, and 100 mM Tris pH 8. Aliquots at various times were quenched with six volumes of buffer containing 8 M urea, 100 mM Tris-Cl, 100 mM boric acid, 100 mM EDTA and analyzed by denaturing PAGE. Gels were stained using SYBR Gold and imaged on an Amersham Typhoon RGB instrument (Cytiva). Bands were quantified with ImageQuant IQTL 8.1 software and then normalized relative to the sequence length. Kinetic plots were non-linearly fitted to the modified first-order rate equation, y = A (1 – e^-*k*x^), where A represents the fraction of active complex, *k* is the first-order rate constant, x is time, and y is the fraction of ligated product in GraphPad Prism 9.

All oligonucleotides used in this work are listed in Supplementary Data Table 5.

## AUTHOR CONTRIBUTIONS

E.M.B, N. R. K., Z.W., and S.D. designed research; E.M.B., N.R.K., and Z.W. performed research; E.M.B., N.R.K., Z.W., and S.D. analyzed the data; E.M.B., N. R. K., and S.D. wrote the paper. E.M.B. and N.R.K. contributed equally to this manuscript.

## DATA AVAILABILITY

Additional supporting data for this article have been included in the Supplemental Material file. The analysis scripts used above are freely available on GitHub (https://github.com/dasguptalab/ligase_ribozyme_population_dynamics) and Zenodo (https://zenodo.org/records/21976743). Sequencing data are available at CurateND (https://doi.org/10.7274/31048348)

## SUPPLEMENTAL MATERIAL

Supplemental material is available for this article.

## ACKNOWLEDGEMENTS

We thank Professors Jack W. Szostak, Michael E. Pfrender, and Dipankar Sen, as well as members of the DasGupta lab (Nicholas J. Colorito, Taylor D. Opolka, Ziyao Zhu, and Dr. Annyesha Biswas) and the Pfrender lab (Bret Coggins, Neil McAdams, and Nitin Vincent) for their valuable feedback on the manuscript. This work was supported by University of Notre Dame startup funds and a National Science Foundation CAREER Award (Award No. MCB-2540950) to S.D., as well as a Chemistry-Biochemistry-Biology Interface (CBBI) Fellowship and a Lucy Family Institute Graduate Scholarship to N.R.K.

## REFERENCES

1. Ameta, S., Winz, M. L., Previti, C., & Jaschke, A. (2014). Next-generation sequencing reveals how RNA catalysts evolve from random space. Nucleic Acids Res, 42(2), 1303–1310. 10.1093/nar/gkt949

2. Asif, M., & Orenstein, Y. (2020). DeepSELEX: inferring DNA-binding preferences from HT-SELEX data using multi-class CNNs. Bioinformatics, 36(Suppl_2), i634–i642. 10.1093/bioinformatics/btaa789

3. Benner, S. A. (2010). Defining life. Astrobiology, 10(10), 1021–1030. 10.1089/ast.2010.0524

4. Bolger, A. M., Lohse, M., & Usadel, B. (2014). Trimmomatic: a flexible trimmer for Illumina sequence data. Bioinformatics, 30(15), 2114–2120. 10.1093/bioinformatics/btu170

5. Chen, X., Li, N., & Ellington, A. D. (2007). Ribozyme catalysis of metabolism in the RNA world. Chem Biodivers, 4(4), 633–655. 10.1002/cbdv.200790055

6. DasGupta, S. (2020). Molecular crowding and RNA catalysis. Org Biomol Chem, 18(39), 7724–7739. 10.1039/d0ob01695k

7. DasGupta, S., Zhang, S., & Szostak, J. W. (2023). Molecular Crowding Facilitates Ribozyme-Catalyzed RNA Assembly. ACS Cent Sci, 9(8), 1670–1678. 10.1021/acscentsci.3c00547

8. Ditzler, M. A., Lange, M. J., Bose, D., Bottoms, C. A., Virkler, K. F., Sawyer, A. W., Whatley, A. S., Spollen, W., Givan, S. A., & Burke, D. H. (2013). High-throughput sequence analysis reveals structural diversity and improved potency among RNA inhibitors of HIV reverse transcriptase. Nucleic Acids Res, 41(3), 1873–1884. 10.1093/nar/gks1190

9. Fu, L., Niu, B., Zhu, Z., Wu, S., & Li, W. (2012). CD-HIT: accelerated for clustering the next-generation sequencing data. Bioinformatics, 28(23), 3150–3152. 10.1093/bioinformatics/bts565

10. Gu, Z. (2022). Complex Heatmap Visualization. iMeta. doi:10.1002/imt2.43.

11. Higgs, P. G., & Muller, U. F. (2025). Principles of selection of ribozymes from random sequence libraries. Journal of the Royal Society Interface, 22(225). ARTN 20240878 10.1098/rsif.2024.0878

12. Hoinka, J., Berezhnoy, A., Dao, P., Sauna, Z. E., Gilboa, E., & Przytycka, T. M. (2015). Large scale analysis of the mutational landscape in HT-SELEX improves aptamer discovery. Nucleic Acids Res, 43(12), 5699–5707. 10.1093/nar/gkv308

13. Holland, J., & Domingo, E. (1998). Origin and evolution of viruses. Virus Genes, 16(1), 13–21. 10.1023/a:1007989407305

14. Jijakli, K., Khraiwesh, B., Fu, W., Luo, L., Alzahmi, A., Koussa, J., Chaiboonchoe, A., Kirmizialtin, S., Yen, L., & Salehi-Ashtiani, K. (2016). The in vitro selection world. Methods, 106, 3–13. 10.1016/j.ymeth.2016.06.003

15. Jimenez, J. I., Xulvi-Brunet, R., Campbell, G. W., Turk-MacLeod, R., & Chen, I. A. (2013). Comprehensive experimental fitness landscape and evolutionary network for small RNA. Proc Natl Acad Sci U S A, 110(37), 14984–14989. 10.1073/pnas.1307604110

16. Kao, C., Rudisser, S., & Zheng, M. (2001). A simple and efficient method to transcribe RNAs with reduced 3’ heterogeneity. Methods, 23(3), 201–205. 10.1006/meth.2000.1131

17. Kobori, S., Nomura, Y., Miu, A., & Yokobayashi, Y. (2015). High-throughput assay and engineering of self-cleaving ribozymes by sequencing. Nucleic Acids Res, 43(13), e85. 10.1093/nar/gkv265

18. Komarova, N., & Kuznetsov, A. (2019). Inside the Black Box: What Makes SELEX Better? Molecules, 24(19). 10.3390/molecules24193598

19. Magoc, T., & Salzberg, S. L. (2011). FLASH: fast length adjustment of short reads to improve genome assemblies. Bioinformatics, 27(21), 2957–2963. 10.1093/bioinformatics/btr507

20. Martin, L. L., Unrau, P. J., & Muller, U. F. (2015). RNA synthesis by in vitro selected ribozymes for recreating an RNA world. Life (Basel), 5(1), 247–268. 10.3390/life5010247

21. Nowosad, J. (2018). CARTOColors’ Palettes. R package version 1.0.0. https://jakubnowosad.com/rcartocolor/.

22. Papastavrou, N., Horning, D. P., & Joyce, G. F. (2024). RNA-catalyzed evolution of catalytic RNA. Proc Natl Acad Sci U S A, 121(11), e2321592121. 10.1073/pnas.2321592121

23. Peri, G., Gibard, C., Shults, N. H., Crossin, K., & Hayden, E. J. (2022). Dynamic RNA Fitness Landscapes of a Group I Ribozyme during Changes to the Experimental Environment. Mol Biol Evol, 39(3). 10.1093/molbev/msab373

24. Pitt, J. N., & Ferre-D’Amare, A. R. (2010). Rapid construction of empirical RNA fitness landscapes. Science, 330(6002), 376–379. 10.1126/science.1192001

25. Pressman, A., Moretti, J. E., Campbell, G. W., Muller, U. F., & Chen, I. A. (2017). Analysis of in vitro evolution reveals the underlying distribution of catalytic activity among random sequences. Nucleic Acids Res, 45(14), 8167–8179. 10.1093/nar/gkx540

26. Pressman, A. D., Liu, Z., Janzen, E., Blanco, C., Muller, U. F., Joyce, G. F., Pascal, R., & Chen, I. A. (2019). Mapping a Systematic Ribozyme Fitness Landscape Reveals a Frustrated Evolutionary Network for Self-Aminoacylating RNA. J Am Chem Soc, 141(15), 6213–6223. 10.1021/jacs.8b13298

27. Robertson, M. P., & Joyce, G. F. (2012). The origins of the RNA world. Cold Spring Harb Perspect Biol, 4(5). 10.1101/cshperspect.a003608

28. Saha, R., Pohorille, A., & Chen, I. A. (2014). Molecular crowding and early evolution. Orig Life Evol Biosph, 44(4), 319–324. 10.1007/s11084-014-9392-3

29. Schlosser, K., & Li, Y. (2005). Diverse evolutionary trajectories characterize a community of RNA-cleaving deoxyribozymes: a case study into the population dynamics of in vitro selection. J Mol Evol, 61(2), 192–206. 10.1007/s00239-004-0346-7

30. Schutze, T., Wilhelm, B., Greiner, N., Braun, H., Peter, F., Morl, M., Erdmann, V. A., Lehrach, H., Konthur, Z., Menger, M., Arndt, P. F., & Glokler, J. (2011). Probing the SELEX process with next-generation sequencing. PLoS One, 6(12), e29604. 10.1371/journal.pone.0029604

31. Spiegelman, S. (1971). An approach to the experimental analysis of precellular evolution. Q Rev Biophys, 4(2), 213–253. 10.1017/s0033583500000639

32. Team, R. C. (2023). A Language and Environment for Statistical Computing. R Foundation for Statistical Computing, Vienna. https://www.R-project.org/.

33. Walton, T., DasGupta, S., Duzdevich, D., Oh, S. S., & Szostak, J. W. (2020). In vitro selection of ribozyme ligases that use prebiotically plausible 2-aminoimidazole-activated substrates. Proc Natl Acad Sci U S A, 117(11), 5741–5748. 10.1073/pnas.1914367117

34. Wickham, H. (2023). stringr: Simple, Consistent Wrappers for Common String Operations. R package version 1.5.1. https://stringr.tidyverse.org.

35. Wickham, H. F., R.; Henry, L.; Müller, K.; Vaughan, D. (2023). dplyr: A Grammar of Data Manipulation. R package version 1.1.4. https://dplyr.tidyverse.org.

36. Wickham, H. P., T.; Seidel, D. (2025). scales: Scale Functions for Visualization. R package version 1.4.0. https://scales.r-lib.org.

37. Wilson, D. S., & Szostak, J. W. (1999). In vitro selection of functional nucleic acids. Annu Rev Biochem, 68, 611–647. 10.1146/annurev.biochem.68.1.611

38. Yang, X., Liu, X., Lou, C., Chen, J., & Ouyang, Q. (2009). A case study of the dynamics of in vitro DNA evolution under constant selection pressure. J Mol Evol, 68(1), 14–27. 10.1007/s00239-008-9182-5

