## Supplementary Information for "Population dynamics of ribozymes during *in vitro* selection"

**Supplementary Data Table 1.** Bioinformatics workflow listing the steps taken to pre-process and analyze the raw sequencing data from the eight rounds of selection (See Supplementary Fig. 1). Each step is numbered and includes a description with its purpose, the method used, the input data, and the output generated. See Methods for a more detailed description.

| Step | Description | Method | Input Data | Output Data |
|---|---|---|---|---|
| **DATA PRE-PROCESSING** | | | | |
| 1 | Merge the paired-end reads for each round (maximum overlap of 81-bp) | FLASH | Raw | Merged reads |
| 2 | Filter reads by quality (Q>30) and remove adapter sequences | Trimmomatic | Step 1 | High quality reads |
| 3 | Filter reads to keep only those that contain the constant 8-bp region flanking the 40-nt variable region | Custom BASH script | Step 2 | Structure filtered reads (Supplementary Data Table 2, Row 4) |
| 4 | Trim reads to retain only the 40-nt variable region | Custom BASH script | Step 3 | Trimmed reads |
| 5 | Combine the merged read files with the unmerged forward read files to retain the most data possible before quality control | Custom BASH script | Step 4 | Combined reads |
| 6 | De-replicate sequences to identify unique sequences and their read counts | Custom BASH script | Step 5 | De-replicated sequences (Supplementary Data Table 2, Row 5) |
| 7 | Filter to keep unique sequences that had >2 reads | Custom BASH script | Step 6 | Filtered sequences |
| 8 | Order the unique sequences according to their read counts | Custom BASH script | Step 7 | Ordered sequences |
| **IDENTIFICATION AND CHARACTERIZATION OF SEQUENCE FAMILIES** | | | | |
| 9 | Cluster round 8 unique sequences according to sequence similarity (sequence identity threshold of 0.9) | CD-HIT-EST | Step 8 | Representative peak sequences |
| 10 | Determine the ten most abundant sequence families in round 8 and identify their peak sequences. Ensure at least 10% dissimilarity between peaks | Custom BASH script | Step 9 | Ten most abundant families and peaks (Supplementary Data Table 3) |
| 11 | Identify unique sequences in each round that had at least 90% identity to any of the round 8 peak sequences | Custom R script | Steps 8 and 10 | Family sequences (Supplementary Figs. 2, 3) |
| **CHARACTERIZATION OF POPULATION DYNAMICS** | | | | |
| 12 | Quantify the abundance of the unique sequences in each round | Custom BASH script | Steps 6 and 8 | Sequence abundances |
| 13 | Determine the ten most abundant unique sequences in each round and determine their similarity to the round 8 families and peak sequences | Custom BASH script | Step 12 | Ten most abundant sequences (Table 1, Fig. 3) |
| **IDENTIFICATION OF CONSERVED FEATURES** | | | | |
| 14 | Calculate the frequency of occurrence for each nucleotide in the most abundant sequence family (Family 1) | Custom R script | Step 10 | Nucleotide conservation of the most abundant sequence family (Fig. 4A) |
| 15 | Identify regions in the unique sequences from each round that are complementary to the substrate | Custom R script | Steps 6 and 8 | Complementary regions to substrate 3' overhang across rounds (Fig. 5) |
| 16 | Identify regions in the families and their peak sequences that are complementary to the substrate in the ten most abundant families | Custom R script | Step 10 | Conserved complementary regions of the families (Supplementary Fig. 6) and peak sequences (Table 2) |
| 17 | Identify regions in the ten most abundant sequences in each round that are complementary to the substrate | Custom R script | Step 13 | Conserved complementary regions of the most abundant sequences (Supplementary Data Table 4) |

**Supplementary Data Table 2.** Progress of selection across eight rounds. Selection pressure was increased across rounds by lowering the concentration of $Mg^{2+}$ (from 20 mM in round 1 to 5 mM in round 8) and reaction times (120 min in round 1 to 10 min in round 8). 582,260-1,067,585 high-quality reads were obtained for rounds 1-8 after quality filtering (See Methods). Sequence diversity decreased from >98% in rounds 1-5 to 54.61% in round 6 and eventually to 12.20% in round 8, indicating significant sequence enrichment. This is consistent with a marked increase in activity of the RNA pool (Fig. 1B). The percent sequence diversity was calculated by dividing the number of unique sequences by the number of high-quality reads for each round.

| | **Round 1** | **Round 2** | **Round 3** | **Round 4** | **Round 5** | **Round 6** | **Round 7** | **Round 8** |
|---|---|---|---|---|---|---|---|---|
| Reaction Time (min) | 120 | 60 | 30 | 20 | 30 | 10 | 10 | 10 |
| $Mg^{2+}$ (mM) | 20 | 20 | 20 | 20 | 20 | 20 | 10 | 5 |
| Total Raw Reads | 1,485,536 | 1,533,916 | 1,649,680 | 1,436,328 | 1,937,410 | 2,336,945 | 1,229,247 | 1,756,169 |
| High Quality Reads (Q>30) | 1,039,660 | 1,067,585 | 1,033,048 | 866,423 | 981,844 | 916,485 | 582,260 | 889,374 |
| Unique Sequences | 1,036,229 | 1,063,996 | 1,029,483 | 863,123 | 966,495 | 500,507 | 92,366 | 108,529 |
| Percent Unique Sequences | 99.67 | 99.66 | 99.65 | 99.62 | 98.44 | 54.61 | 15.86 | 12.20 |

**Supplementary Data Table 3.** The ten most abundant ribozyme families are shown along with their relative abundances and most abundant (peak) sequences (See Table 2). Each sequence corresponds to the 40-nucleotide variable region of the most abundant member (peak sequence) within the family. The percent abundance corresponds to the total percent abundance of that family, which was calculated as the sum of all read counts in a family divided by the total read counts in round 8. The percent peak abundance is calculated as the number of sequence reads corresponding to the peak divided by the total number of sequence reads in the family. See Fig. 1C for an illustration of how each family was enriched during selection.

| Family | Number of Sequences | Read Counts | Percent Abundance | Peak Sequence (40-nt variable region; 5'→3') | Percent Peak Abundance |
|---|---|---|---|---|---|
| 1 | 1460 | 324773 | 36.52 | GAATGCTGCCAACCGTGCGGGCTAATTGGCAGACTGAGCT | 79.01 |
| 2 | 264 | 91636 | 10.30 | CCTAGCTAGCGCTGACTAGGACAGATGAGCGGCGGAACCA | 89.57 |
| 3 | 176 | 47376 | 5.33 | TTAGTGAAATTGGTGCCCAAGCAGAGAATTGGGATAAATC | 86.54 |
| 4 | 141 | 37250 | 4.19 | GAACCCTTATCACAGTCGTGCGGATTTGTAAGCCTAAGCG | 91.59 |
| 5 | 128 | 29264 | 3.29 | AGAGACCGTGAGCTTGCGGAATGTTAGCAGAACAGAACTG | 90.03 |
| 6 | 113 | 17774 | 1.99 | AGCCACTGCGGAAGACCTTAAGAGGTGTAATTGCTCACCC | 89.57 |
| 7 | 112 | 16553 | 1.86 | AAAAGTTTCGCTGAATTGGACAGACCACCGCGTGAAGTGG | 87.23 |
| 8 | 87 | 9579 | 1.08 | CCGGTCTTAAGCCCCTGCGTTGCGGAAATGCACGTTGCCC | 88.25 |
| 9 | 87 | 7336 | 0.82 | GACGGACGGTCGCGGTAACCTATGACCATGAGACGGAACA | 89.35 |
| 10 | 67 | 4539 | 0.51 | TGATGAATCGGCATACGTGGGTCAGAGTCATAGTGCGACA | 89.07 |

**Supplementary Data Table 4.** Selection dynamics of base-pair complementarity between ligase ribozyme sequences and the substrate. All nineteen sequences that comprise the top ten sequences in each round (See Table 1) exhibit extensive base-pair complementarity (BPC) with the substrate 3′ overhang sequence. Ribozyme nucleotides that have Watson-Crick complementarity with the substrate are indicated in red, and wobble complementarity is indicated in blue. The nucleotides contained in the sliding window are underlined in each sequence (See Supplementary Fig. 5). The substrate 3′ overhang is underlined. Complementarity is denoted as in Table 2.

| Seq. ID | Sequence Rank in each round | | | | | | | | Substrate (5′→3′): ACCACCGCAUUCCGCA | | |
|---|---|---|---|---|---|---|---|---|---|---|---|
| | 1 | 2 | 3 | 4 | 5 | 6 | 7 | 8 | Sequence (40-nt variable region; 5′→3′) | Family | BPC |
| | | | | | | | | | 1 5 10 15 20 25 30 35 40 | | |
| 1_1 | 1 | 1 | 1 | 1 | 2 | 1 | 1 | 1 | GAAUGCUGCCAACCGUGCGGGCUAAUUGGCAGACUGAGCU | 1 | 6/8 |
| 1_2 | 2 | 2 | 3 | 2 | 1 | 2 | 2 | 3 | UUAGUGAAAUUGGUGCCCAAGCAGAGAAUUGGGAUAAAUC | 3 | 6/8 |
| 1_3 | 3 | 3 | 2 | 3 | 4 | 6 | 4 | 2 | CCUAGCUAGCGCUGACUAGGACAGAUGAGCGGCGGAACCA | 2 | 6/8 |
| 1_4 | 4 | 5 | 7 | 9 | 10 | 9 | 7 | 4 | GAACCCUUAUCACAGUCGUGCGGAUUUGUAAGCCUAAGCG | 4 | 6/8 |
| 1_5 | 5 | 4 | 5 | 6 | 5 | 3 | 3 | 9 | CUGGCAAACACAGCGCGCUGUGUGUUAUGUGGGCGGUCU | - | 7/8 |
| 1_6 | 6 | 7 | 4 | 7 | 6 | 4 | 6 | 10 | UCAGUCGGAGUACCAGAGCGAUAGACGUCCCCGGAAGCCG | - | 6/8 |
| 1_7 | 7 | 8 | 8 | 4 | - | - | 9 | 5 | AGAGACCGUGAGCUUGCGGAAUGUUAGCAGAACAGAACUG | 5 | 8/8 |
| 1_8 | 8 | 6 | 10 | 5 | 3 | 5 | 5 | - | UUGGUGUAGAGCGCCAACUGGACAGACCUUACGGAAACGG | - | (3_3)/8 |
| 1_9 | 9 | - | - | - | 7 | - | 8 | - | GACGGACGGUCGCGGUAACCUAUGACCAUGAGACGGAACA | 9 | 5/8 |
| 1_10 | 10 | - | - | - | - | 7 | 10 | - | UGUCGUUGAGAUAUACUGGACAGACAAGACGUGGGAACUG | - | 6/8 |
| 2_9 | - | 9 | - | - | - | - | - | - | UGAUGAAUCGGCAUACGUGGGUCAGAGUCAUAGUGCGACA | 10 | 5/8 |
| 2_10 | - | 10 | - | - | 9 | - | - | - | UGAUCGGCAACCGUGGUAUAAGUUCACAAUGUGCGGCAGG | - | 5/8 |
| 3_6 | - | - | 6 | 8 | - | - | - | - | UUAGAGAGCCACAUGCGCUCUGUUUUGCGGAUAAAAUGUG | - | 6/8 |
| 3_9 | - | - | 9 | - | - | - | - | - | AAUUACCUGCCCAUGUGCUGAAUGCAGCGAAAUCAUCGAA | - | (3_4)/8 |
| 4_10 | - | - | - | 10 | - | - | - | 6 | AGCCACUGCGGAAGACCUUAAGAGGUGUAAUUGCUCACCC | 6 | 7/8 |
| 5_8 | - | - | - | - | 8 | 8 | - | - | CGCAGCGCUCACCUUGAGAGGUCAGAAAAGAUGUUGAAUA | - | (3_4)/8 |
| 6_10 | - | - | - | - | - | 10 | - | - | GAAUGCGUCCACUUGCGGGCACAAACUCGACGACUGAGCA | - | 6/8 |
| 8_7 | - | - | - | - | - | - | - | 7 | AAAAGUUUCGCUGAAUUGGACAGACCACCGCGUGAAGUGG | 7 | (3_3)/8 |
| 8_8 | - | - | - | - | - | - | - | 8 | GAAUGCUACCAACCGUGCGGGCUAAUUGGCAGACUGAGCU | - | 6/8 |
| | | | | | | | | | 1 5 10 15 20 25 30 35 40 | | |

**Supplementary Data Table 5.** Oligonucleotides used in this work. Nucleotides in the 40-nt variable region in the ligase ribozyme sequences are shown in boldface, mutations are shown in red, the $U_6$ linker is shown in italics, and the primer sequence is underlined. The T7 promoter sequence in the forward primer PCR_Fwd_primer is highlighted in purple. The first two residues in PCR_Rvs_primer are 2′-*O*-methyl modified, to prevent 3′ heterogeneity in runoff transcription, which is indicated by ‘m’. Oligonucleotides were either purchased from Integrated DNA Technologies (IDT) or generated enzymatically by *in vitro* transcription (IVT) of dsDNA templates. AI-Lig was generated by incubating P-Lig with EDC and 2-aminoimidazole (2AI) (See ‘RNA preparation and ligation assays’ in Materials and Methods).

| No. | Oligo | Sequence (5→′3′) | Type | Source |
|---|---|---|---|---|
| 1 | F1 Peak (Seq1_1) | GACUCACUGACACAGAUCCACUCACGGACAGCG**GAAUGCUGCCAACCG**<br>**UGCGGGCUAAUUGGCAGACUGAGCU**CGCUGUCC*UUUUUU*GGCUAAGG | RNA | IVT |
| 2 | F1 Peak _U4A | GACUCACUGACACAGAUCCACUCACGGACAGCG**GAAAGCUGCCAACCG**<br>**UGCGGGCUAAUUGGCAGACUGAGCU**CGCUGUCC*UUUUUU*GGCUAAGG | RNA | IVT |
| 3 | F1 Peak _U4C | GACUCACUGACACAGAUCCACUCACGGACAGCG**GAACGCUGCCAACCG**<br>**UGCGGGCUAAUUGGCAGACUGAGCU**CGCUGUCC*UUUUUU*GGCUAAGG | RNA | IVT |
| 4 | F1 Peak _U4G | GACUCACUGACACAGAUCCACUCACGGACAGCG**GAAGGCUGCCAACCG**<br>**UGCGGGCUAAUUGGCAGACUGAGCU**CGCUGUCC*UUUUUU*GGCUAAGG | RNA | IVT |
| 5 | F1 Peak _G8A | GACUCACUGACACAGAUCCACUCACGGACAGCG**GAAUGCUACCAACCG**<br>**UGCGGGCUAAUUGGCAGACUGAGCU**CGCUGUCC*UUUUUU*GGCUAAGG | RNA | IVT |
| 6 | F1 Peak _G8C | GACUCACUGACACAGAUCCACUCACGGACAGCG**GAAUGCUCCCAACCG**<br>**UGCGGGCUAAUUGGCAGACUGAGCU**CGCUGUCC*UUUUUU*GGCUAAGG | RNA | IVT |
| 7 | F1 Peak _G8U | GACUCACUGACACAGAUCCACUCACGGACAGCG**GAAUGCUUCCAACCG**<br>**UGCGGGCUAAUUGGCAGACUGAGCUCGCU**GUCC*UUUUUU*GGCUAAGG | RNA | IVT |
| 8 | F1 Peak _G17A | GACUCACUGACACAGAUCCACUCACGGACAGCG**GAAUGCUGCCAACCG**<br>**UACGGGCUAAUUGGCAGACUGAGCU**CGCUGUCC*UUUUUU*GGCUAAGG | RNA | IVT |
| 9 | F1 Peak _G17C | GACUCACUGACACAGAUCCACUCACGGACAGCG**GAAUGCUGCCAACCG**<br>**UCCGGGCUAAUUGGCAGACUGAGCU**CGCUGUCC*UUUUUU*GGCUAAGG | RNA | IVT |
| 10 | F1 Peak _G17U | GACUCACUGACACAGAUCCACUCACGGACAGCG**GAAUGCUGCCAACCG**<br>**UUCGGGCUAAUUGGCAGACUGAGCU**CGCUGUCC*UUUUUU*GGCUAAGG | RNA | IVT |
| 11 | F1 Peak _G20A | GACUCACUGACACAGAUCCACUCACGGACAGCG**GAAUGCUGCCAACCG**<br>**UGCGAGCUAAUUGGCAGACUGAGCU**CGCUGUCC*UUUUUU*GGCUAAGG | RNA | IVT |
| 12 | F1 Peak _G20C | GACUCACUGACACAGAUCCACUCACGGACAGCG**GAAUGCUGCCAACCG**<br>**UGCGCGCUAAUUGGCAGACUGAGCU**CGCUGUCC*UUUUUU*GGCUAAGG | RNA | IVT |
| 13 | F1 Peak _G20U | GACUCACUGACACAGAUCCACUCACGGACAGCG**GAAUGCUGCCAACCG**<br>**UGCGUGCUAAUUGGCAGACUGAGCU**CGCUGUCC*UUUUUU*GGCUAAGG | RNA | IVT |
| 14 | F1 Peak _A37C | GACUCACUGACACAGAUCCACUCACGGACAGCG**GAAUGCUGCCAACCG**<br>**UGCGGGCUAAUUGGCAGACUGCGCU**CGCUGUCC*UUUUUU*GGCUAAGG | RNA | IVT |
| 15 | F1 Peak _A37G | GACUCACUGACACAGAUCCACUCACGGACAGCG**GAAUGCUGCCAACCG**<br>**UGCGGGCUAAUUGGCAGACUGGGCU**CGCUGUCC*UUUUUU*GGCUAAGG | RNA | IVT |
| 16 | F1 Peak _A37U | GACUCACUGACACAGAUCCACUCACGGACAGCG**GAAUGCUGCCAACCG**<br>**UGCGGGCUAAUUGGCAGACUGUGCU**CGCUGUCC*UUUUUU*GGCUAAGG | RNA | IVT |
| 17 | Seq1_2 | GACUCACUGACACAGAUCCACUCACGGACAGCG**UUAGUGAAAUUGGUG**<br>**CCCAAGCAGAGAAUUGGGAUAAAUC**CGCUGUCC*UUUUUU*GGCUAAGG | RNA | IVT |
| 18 | Seq1_3 | GACUCACUGACACAGAUCCACUCACGGACAGCG**CCUAGCUAGCGCUGA**<br>**CUAGGACAGAUGAGCGGCGGAACCA**CGCUGUCC*UUUUUU*GGCUAAGG | RNA | IVT |
| 19 | Seq1_4 | GACUCACUGACACAGAUCCACUCACGGACAGCG**GAACCCUUAUCACAG**<br>**UCGUGCGGAUUUGUAAGCCUAAGCG**CGCUGUCC*UUUUUU*GGCUAAGG | RNA | IVT |
| 20 | Seq1_5 | GACUCACUGACACAGAUCCACUCACGGACAGCG**CUGGCAAACACAGCG**<br>**CGCUGUGUGUUAUGUGGGGCGGUCU**CGCUGUCC*UUUUUU*GGCUAAGG | RNA | IVT |

| | | | | |
|---|---|---|---|---|
| 21 | Seq1_6 | GACUCACUGACACAGAUCCACUCACGGACAGCG**UCAGUCGGAGUACCA GAGCGAUAGACGUCCCCGGAAGCCG**CGCUGUCC*UUUUUU*GGCUAAGG | RNA | IVT |
| 22 | Seq1_7 | GACUCACUGACACAGAUCCACUCACGGACAGCG**AGAGACCGUGAGCUU GCGGAAUGUUAGCAGAACAGAACUG**CGCUGUCC*UUUUUU*GGCUAAGG | RNA | IVT |
| 23 | Seq1_8 | GACUCACUGACACAGAUCCACUCACGGACAGCG**UUGGUGUAGAGCGCC AACUGGACAGACCUUACGGAAACGG**CGCUGUCC*UUUUUU*GGCUAAGG | RNA | IVT |
| 24 | Seq1_9 | GACUCACUGACACAGAUCCACUCACGGACAGCG**GACGGACGGUCGCGG UAACCUAUGACCAUGAGACGGAACA**CGCUGUCC*UUUUUU*GGCUAAGG | RNA | IVT |
| 25 | Seq1_10 | GACUCACUGACACAGAUCCACUCACGGACAGCG**UGUCGUUGAGAUAUA CUGGACAGACAAGACGUGGGAACUG**CGCUGUCC*UUUUUU*GGCUAAGG | RNA | IVT |
| 26 | Seq2_9 | GACUCACUGACACAGAUCCACUCACGGACAGCG**UGAUGAAUCGGCAUA CGUGGGUCAGAGUCAUAGUGCGACA**CGCUGUCC*UUUUUU*GGCUAAGG | RNA | IVT |
| 27 | Seq2_10 | GACUCACUGACACAGAUCCACUCACGGACAGCG**UGAUCGGCAACCGUG GUAUAAGUUCACAAUGUGCGGCAGG**CGCUGUCC*UUUUUU*GGCUAAGG | RNA | IVT |
| 28 | Seq3_6 | GACUCACUGACACAGAUCCACUCACGGACAGCG**UUAGAGAGCCACAUG CGCUCUGUUUUGCGGAUAAAAUGUG**CGCUGUCC*UUUUUU*GGCUAAGG | RNA | IVT |
| 29 | Seq3_9 | GACUCACUGACACAGAUCCACUCACGGACAGCG**AAUUACCUGCCCAUG UGCUGAAUGCAGCGAAAUCAUCGAA**CGCUGUCC*UUUUUU*GGCUAAGG | RNA | IVT |
| 30 | Seq4_10 | GACUCACUGACACAGAUCCACUCACGGACAGCG**AGCCACUGCGGAAGA CCUUAAGAGGUGUAAUUGCUCACCC**CGCUGUCC*UUUUUU*GGCUAAGG | RNA | IVT |
| 31 | Seq5_8 | GACUCACUGACACAGAUCCACUCACGGACAGCG**CGCAGCGCUCACCUU GAGAGGUCAGAAAAGAUGUUGAAUA**CGCUGUCC*UUUUUU*GGCUAAGG | RNA | IVT |
| 32 | Seq6_10 | GACUCACUGACACAGAUCCACUCACGGACAGCG**GAAUGCGUCCACUUG CGGGCACAAACUCGACGACUGAGCA**CGCUGUCC*UUUUUU*GGCUAAGG | RNA | IVT |
| 33 | Seq8_7 | GACUCACUGACACAGAUCCACUCACGGACAGCG**AAAAGUUUCGCUGAA UUGGACAGACCACCGCGUGAAGUGG**CGCUGUCC*UUUUUU*GGCUAAGG | RNA | IVT |
| 34 | Seq8_8 | GACUCACUGACACAGAUCCACUCACGGACAGCG**GAAUGCUACCAACCG UGCGGGCUAAUUGGCAGACUGAGCU**CGCUGUCC*UUUUUU*GGCUAAGG | RNA | IVT |
| 35 | P-Lig | (5’-monophosphate) – ACCACCGCAUUCCGCA | RNA | IDT |
| 36 | AI-Lig | (5’-phosphoro-2-aminoimidazole) – ACCACCGCAUUCCGCA | RNA | Activation of P-Lig |
| 37 | AI-Lig-Biotin | (5'-phosphoro-2-aminoimidazole) – ACCACCGCAUUCCGCA -(dC-biotin) | RNA | Biotinylat-ion of AI-Lig |
| 38 | Template | GCGGUGGUCCUUAGCC | RNA | IDT |
| 39 | RQ1 | GTGCGGAATGCGGTGGT | DNA | IDT |
| 40 | RQ2 | GCCTTAGCCAAAAAAGGACAGCG | DNA | IDT |
| 41 | SeqPCR1 | GTTCAGAGTTCTACAGTCCGACGATCCGGTAGGTCCCTTAGCCAAAAA AGGACAGCG | DNA | IDT |
| 42 | SeqPCR2 | AGACGTGTGCTCTTCCGATCTGACTCACTGACACAGATCCACTCAC | DNA | IDT |
| 43 | SeqPCR3 | GTTCAGAGTTCTACAGTCCGACGATC | DNA | IDT |
| 44 | SeqPCR4 | AGACGTGTGCTCTTCCGATCT | DNA | IDT |
| 45 | RT_Primer | GTGCGGAATGCGGTGGTCCTT | DNA | IDT |
| 46 | PCR_Fwd primer | TAATACGACTCACTATAGACTCACTGACAC | DNA | IDT |
| 47 | PCR_Rvs primer | mCmCTTAGCCAAAAAAGGACAGCG | DNA | IDT |

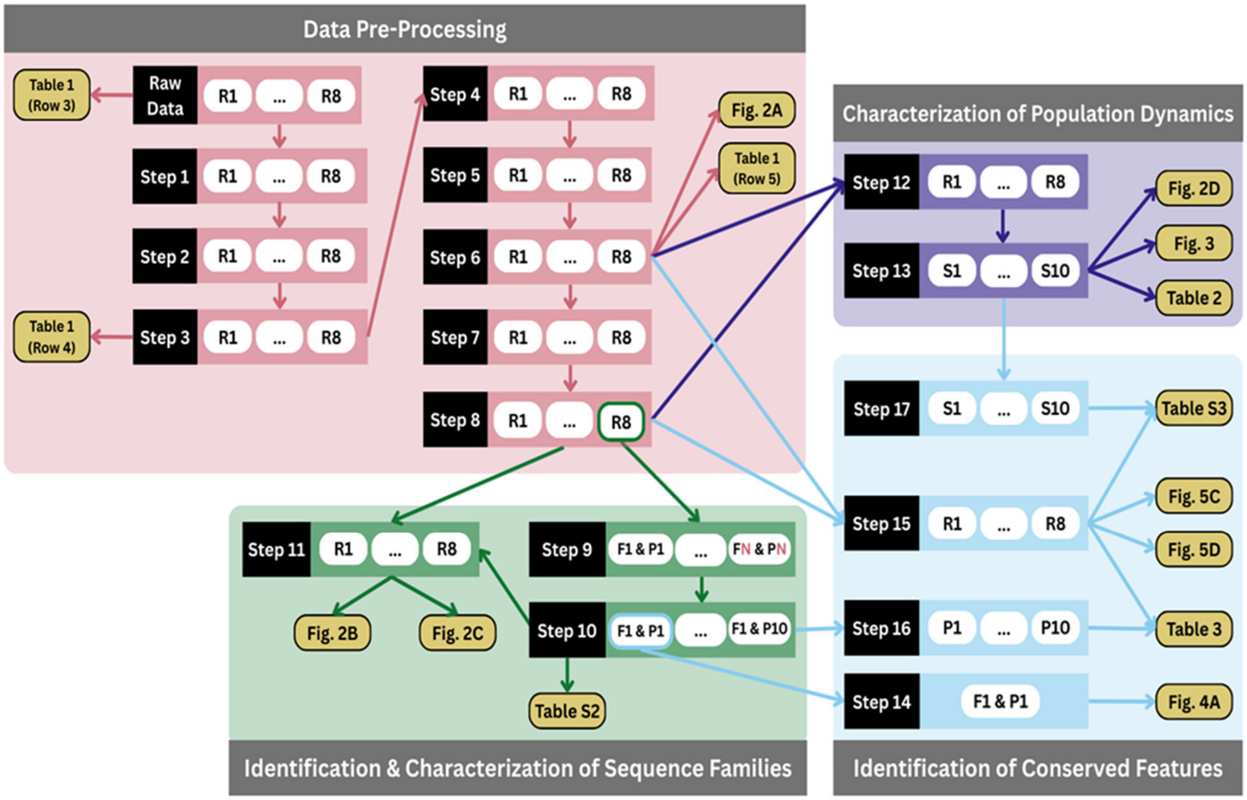


**Supplementary Fig. 1.** The bioinformatics analysis workflow entails several steps (see Supplementary Table 1). The workflow begins with pre-processing raw sequencing data in steps 1 through 8 (pink box). Sequence families and representative peak sequences were identified in steps 9 through 11 (green box). Population dynamics of the ribozyme sequences were characterized in steps 12 and 13 (purple box). Nucleotide conservation and regions complementary to the substrate 3′ overhang were identified in steps 14 through 17 (blue box). The tables and figures produced in these various stages of bioinformatics analysis are shown in yellow.

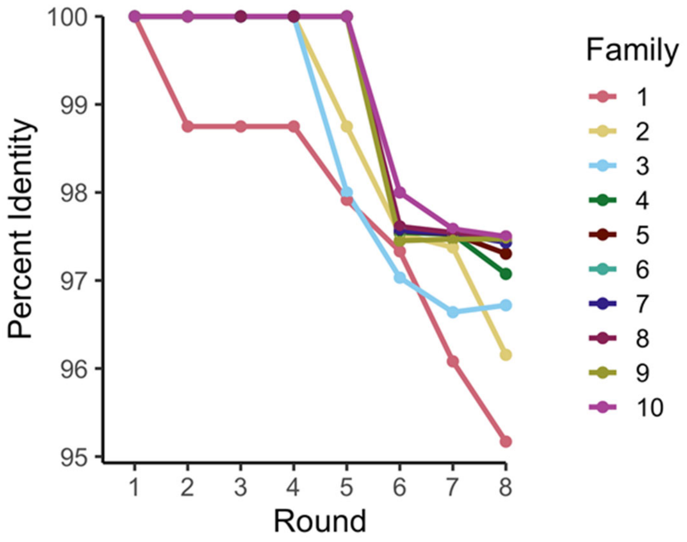


**Supplementary Fig. 2.** Sequence diversity within the ten most abundant families increased during selection, due to the creation of sequence variants of the peak sequence in each family. The sequence diversity within each family increased with sequence family abundance.

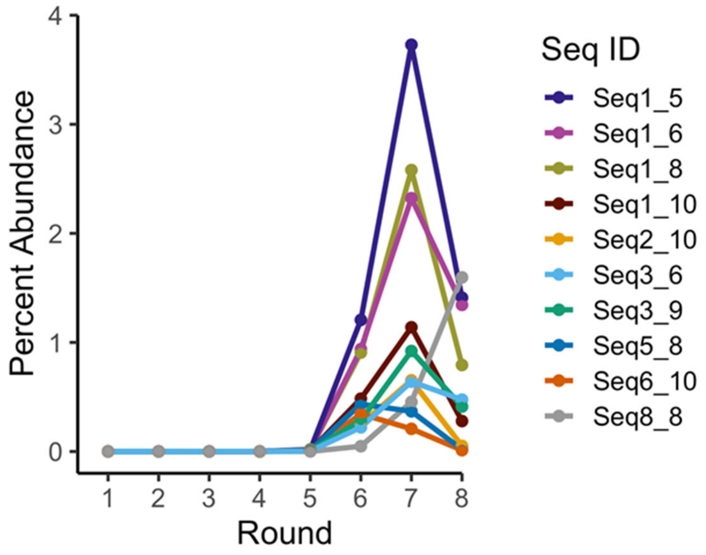


**Supplementary Fig. 3.** Enrichment patterns of non-peak sequences that emerged in the top ten in any of the eight selection rounds (Supplementary Data Table 4). The percent abundances in each round were calculated by dividing the number of reads of each sequence or family by the total number of reads in each round.

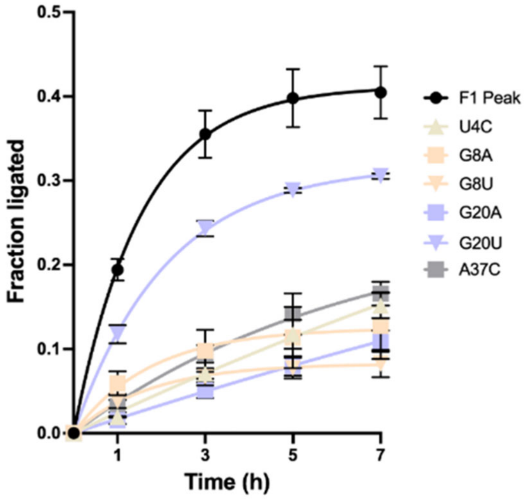


**Supplementary Fig. 4.** Kinetic profiles for RNA ligation catalyzed by single point mutants of the F1 peak sequence identified through nucleotide conservation analysis in Fig. 4A. Point mutations to the F1 peak sequence reduce activity to different extents (see Fig. 4E for gel data and Fig. 4F for $k_{obs}$ values). Error bars indicate standard deviation. All data were obtained from triplicate measurements. Ligation reactions contained 1 μM ribozyme, 1.2 μM RNA template, and 2 μM RNA substrate in 100 mM Tris-HCl (pH 8.0), 300 mM NaCl, and 10 mM $MgCl_2$.

#

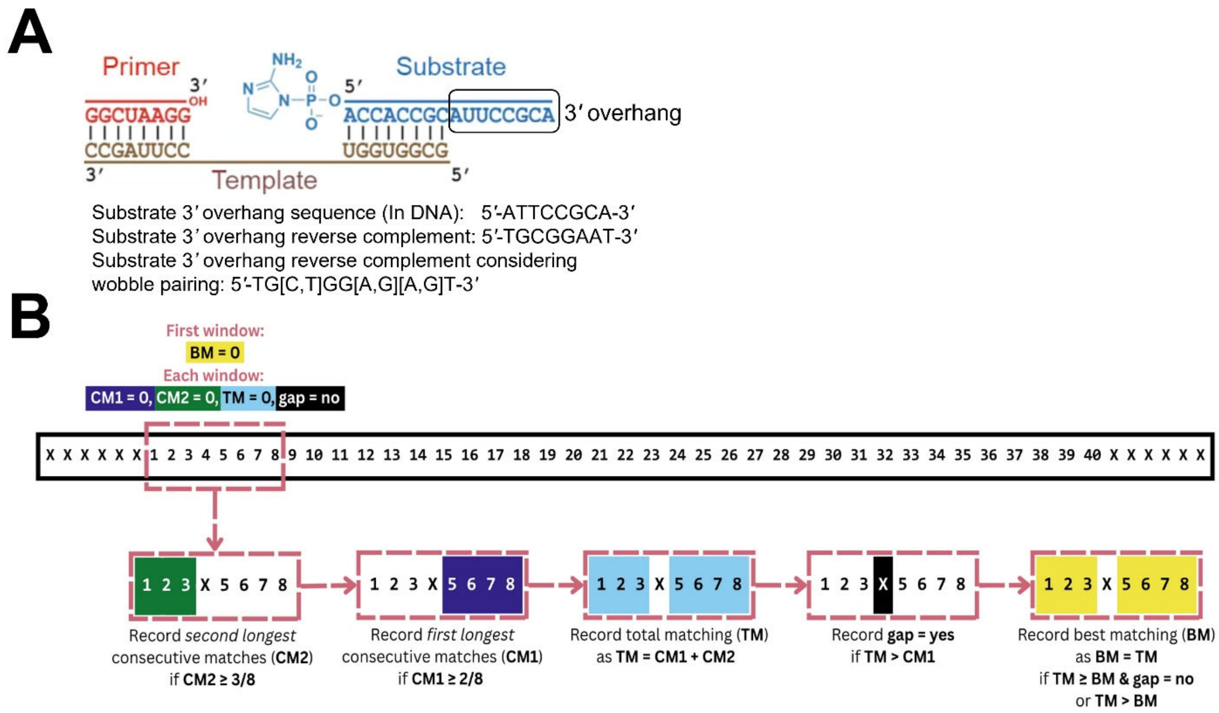


**Supplementary Fig. 5.** Custom algorithm for identifying regions in the ribozyme sequences that are complementary to the substrate. **(A)** Ligation junction showing base-pairing between the 16-nt template and both the 3′ end of the ribozyme (primer) and the first 8 nucleotides of the 2AI-activated substrate (AI-Lig). The last 8 nucleotides of AI-Lig (substrate 3′ overhang) do not base-pair with the template and are available to pair with the ribozyme. **(B)** We used an 8-nt sliding window to identify sequence regions complementary to the substrate 3’ overhang. Our algorithm includes wobble pairs in the search and allows for up to two mismatches. For the first sliding window, the number of best matching (BM) nucleotides was set to zero. For each 8-nt window, the remaining variables were reset to zero. Then the first longest stretch of consecutive matches (CM1) was counted, recording the number of matching nucleotides only when CM1 ≥ 2 nucleotides. Therefore, this requires a minimum of a 2-nt complementarity. This was followed by counting the second longest stretch of consecutive matches (CM2), recording the number of matching nucleotides only if CM2 ≥3 nucleotides, thus, ensuring a minimum 3-nt complementarity on either side of a mismatch. The number of total matching (TM) nucleotides was recorded as CM1 + CM2. A gap was recorded if TM >CM1. Our final step for each sliding window was to record BM as TM if TM ≥BM and there was no gap, or if TM >BM. The requirements to record BM prioritized non-gapped sequences that did not contain internal mismatches. After the last window was processed, BM contained the last best matching complementary region in the sequence.

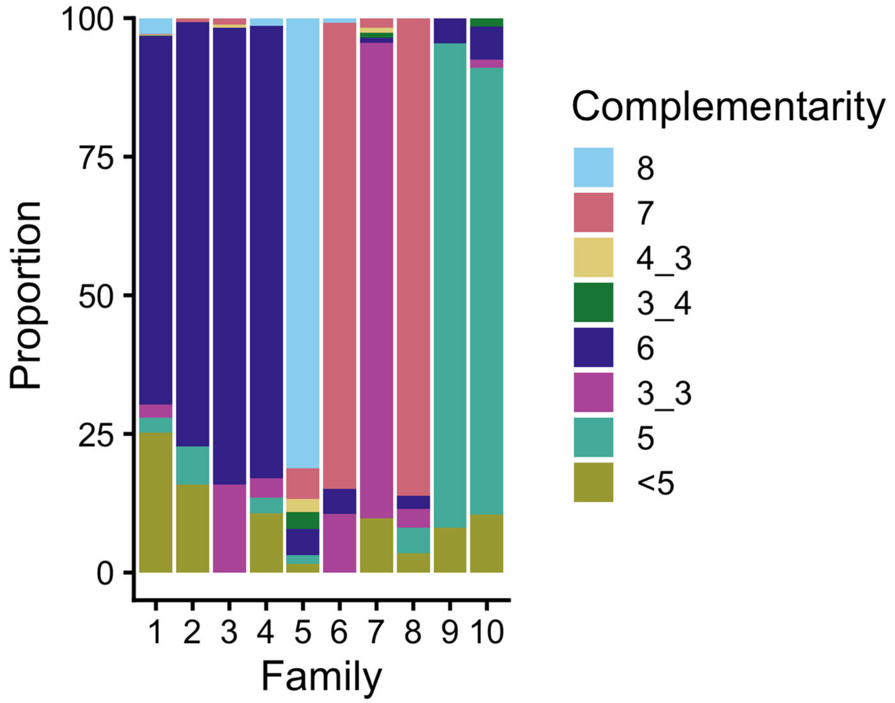


**Supplementary Fig. 6.** Base-pair complementarity for each of the ten most abundant families in round 8. The proportion of complementary nucleotides in the unique sequences (with >2 reads) for each sequence family in round 8. The majority of sequences in each family share the same number of complementary bases with the substrate as the representative peak sequences for that family (see Table 2). Complementarity of 6 base-pairs was most common in the top four most abundant sequence families. The most abundant sequence family (family 1) had the greatest number of sequences with less than 5 base-pair complementarity. Sequence family 5 contained the most sequences with 8 base-pair complementarity. Sequences with internal mismatches that create gaps are denoted with an underscore (e.g., 3_3 indicates two stretches of 3-nt complementarity separated by a mismatch).
